# Ongoing thoughts at rest reflect functional brain organization and behavior

**DOI:** 10.1101/2025.08.18.670664

**Authors:** Jin Ke, Taylor A. Chamberlain, Hayoung Song, Anna Corriveau, Ziwei Zhang, Taysha Martinez, Laura Sams, Marvin M. Chun, Yuan Chang Leong, Monica D. Rosenberg

**Author notes:** Correspondence to: Jin Ke and Monica D. Rosenberg.

## Abstract

Resting-state functional connectivity (rsFC)—brain connectivity observed when people rest with no external tasks—predicts individual differences in behavior. Yet, rest is not idle; it involves streams of thoughts. Are these ongoing thoughts reflected in FC and do they contribute to the relationship between rsFC and behavior? To test this question, we developed an annotated rest paradigm where participants rated and verbally described their thoughts after each rest period during functional MRI (*N* = 60). Our findings revealed rich and idiosyncratic thoughts across individuals. Similarity in thoughts was associated with more similar FC patterns within and across individuals. In addition, both thought ratings and topics could be decoded from FC. Furthermore, neuromarkers of these thoughts generalized to unseen individuals in the Human Connectome Project dataset (*N* = 908), where decoded thought patterns during rest predicted positive vs. negative trait-level individual differences. Together, our findings reveal that ongoing thoughts at rest are reflected in brain dynamics and these network patterns predict everyday cognition and experiences. Understanding subjective in-scanner experiences is thus crucial in characterizing the relationship between individual differences in functional brain organization and behavior.

## Introduction

The human mind wanders frequently. From moment to moment, it continuously generates streams of thoughts that drift from the present to the far reaches of time, and from one topic to another. This pervasive tendency, often referred to as “mind-wandering” or “off-task thoughts” during tasks and spontaneous thinking in the absence of task demands, is not an exception but rather the brain’s default state^1–3^.

Experimental paradigms in cognitive neuroscience have typically viewed the wandering mind as experimental noise. Resting periods, in which participants do not actively engage in any explicit task, were used as a baseline condition under the assumption that any mental activity during these periods constitutes such noise^4^. However, over the past decade neuroimaging studies have revealed that resting-state data contains rich information about cognition and behavior. One fruitful line of research demonstrates that resting-state functional connectivity (rsFC) predicts individual differences in out-of-scanner subject-specific measures, including psychometric performance^5–7^, traits^8–10^, and clinical symptoms^11–14^. Adding to these findings, Smith and colleagues identified a strong mode of brain-behavior association, linking rsFC patterns with an extensive set of behavioral measures in an integrated, multivariate manner^15^. One prevalent hypothesis of rsFC-behavior associations is that individual functional connectome “fingerprints” reflect intrinsic variations in brain architecture^5^ that are causally related to individual differences in behavior.

Prior work has largely disregarded the possibility that rsFC patterns are shaped by individuals’ ongoing mental experiences and thus relate with behavior. Yet, two lines of evidence support such a possibility. First, during resting-state fMRI scans, even in the absence of experimentally imposed external demands or sensory input, the mind remains active^16^. Rich and individualized ongoing experiences during rest^17–25^ may shape rsFC patterns. Second, mind wandering content, including personal goals and planning, concerns, memory, and feelings^3,26–31^, reveals individual internal states and predicts cognitive traits and psychiatric symptoms^32–35^. RsFC may therefore predict behavior in part because individuals with different characteristics engage in different patterns of thinking during rest.

Indeed, recent pioneering studies have begun to provide empirical evidence on the association between rsFC and ongoing cognition^36–39^. In particular, Gonzalez-Castillo and colleagues^37^ used retrospective thought probes and showed that specific aspects of these thoughts can be successfully predicted from rsFC. In addition, Hardikar and colleagues^39^ revealed that individual differences in ongoing thoughts and dispositional traits are related to large-scale cortical gradients—continuous axes of functional connectivity spanning sensory–motor to transmodal cortex. While this work has advanced our understanding of subjective in-scanner experiences and their neural substrates, several critical questions remain unresolved. First, most prior work linked static rsFC estimates with a single snapshot of thoughts, leaving unclear how moment-to-moment fluctuations in these experiences map onto evolving brain dynamics. Addressing this issue is essential to determine whether rsFC reflects active cognitive processes rather than aggregated, trait-like signatures^40–42^.

Second, although rsFC has been linked to ongoing thoughts, it remains untested whether the neural signatures of these thoughts are directly related with external behavior, which is critical to establish that ongoing thoughts contribute to rsFC-behavior associations. Third, prior predictive models of thoughts reported substantial individual variability in the neural representations of mind-wandering^38^, while others provided initial evidence for cross-sample generalization for specific affective dimensions of spontaneous thoughts^24^. These findings raise important questions about the extent to which thought-related connectivity patterns reflect idiosyncratic states rather than reproducible neural signatures. This current study aims to address these gaps in understanding how ongoing thought dynamics relate to brain dynamics and contribute to rsFC-behavior associations. We hypothesize that ongoing thoughts at rest meaningfully vary between individuals, are reflected in ongoing brain dynamics, and thus contribute to the rsFC-behavior associations across datasets.

To test the hypothesis, we were inspired by earlier thought-probing paradigms^20,21,36^ and designed an annotated rest task (**Fig. 1**) as part of a two-session functional magnetic resonance imaging (fMRI) study (*N* = 60) collecting rest, movie-watching, attentional performance, and post-scan memory and engagement in the movies. We named this the Chicago Attention and Thoughts (CAT) dataset. In the annotated rest task, we regularly sampled individuals’ ongoing thoughts between rest periods with subjective ratings and free speech as in-the-moment probes, which enabled us to capture the dynamic nature of ongoing thoughts and allowed for a granular mapping between thoughts and large-scale brain networks^4,16,43^. Moreover, we directly asked participants *what* they were thinking about, expanding the multidimensional experience sampling of ratings on thought dimensions^22,27,44^ to also probe thought semantics. This design allowed us to capture multiple aspects of thought: self-reported dimensions (thought dimensions), semantic content derived from natural language embeddings (thought content), and human-annotated topics (thought topics; **Fig. 1**).

**Figure 1.**
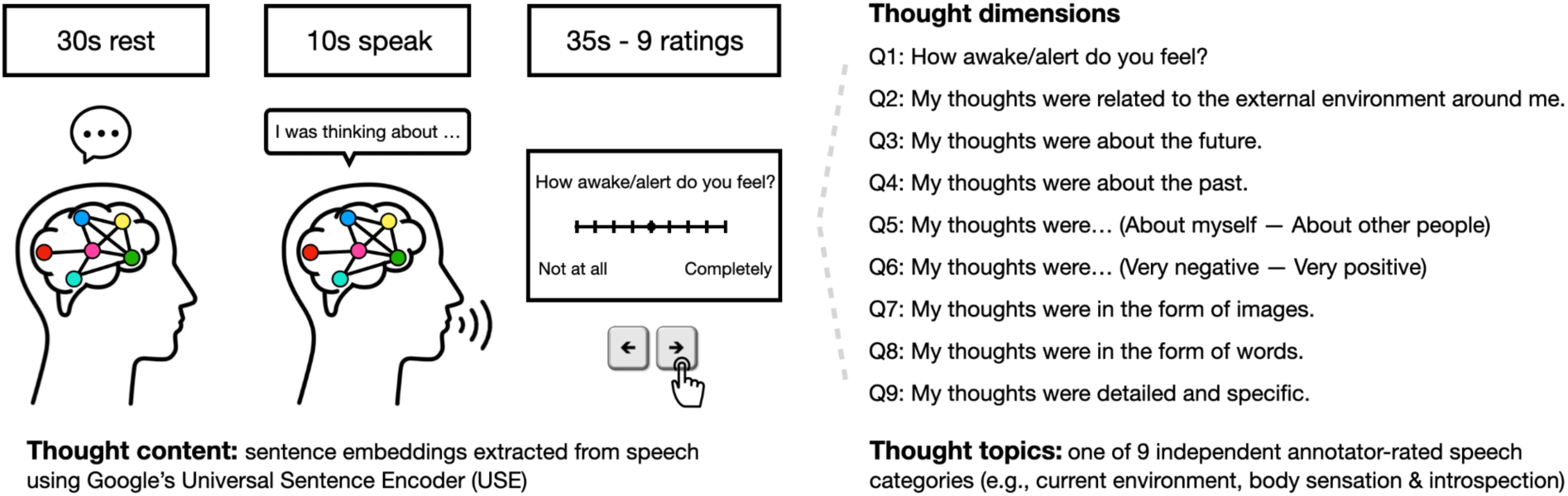
Annotated rest task and experiment design. 60 individuals performed 4 10-minute runs of annotated rest (2 runs per MRI session). Each run had 8 trials, one shown here: participants rested for 30 seconds, verbally reported their ongoing thoughts during rest for 10 seconds, and rated their thoughts on 9 dimensions. Three measures of thoughts were used in the current work: (1) dimensions: the 9 thought dimensions rated by participants in each trial; (2) content: semantic embeddings as 512-dimensional vectors extracted from the transcriptions of participants’ speech using Google’s Universal Sentence Encoder (USE); (3) topics: one of nine topics rated by 5 independent annotators from the transcriptions of participants’ speech. See Fig. 2e for details and **Suppl. Table 2** for definitions of the nine topics.

Using these data, we observed rich and individualized thoughts during resting-state fMRI scans. Representational similarity analysis indicated that individuals with more similar thoughts exhibited more similar rsFC profiles, and that moment-to-moment thought fluctuations tracked rsFC dynamics. Connectome-based models predicted thought dimensions and classified thought topics. These models successfully distinguished an individual with aphantasia—who lacks visual imagery—from their otherwise identical, healthy twin. In addition, these models generalized beyond self-report, predicting non-introspective measures such as pupil size, linguistic sentiment of speech, and strength of a sustained attention network^45,46^. Critically, neuromarkers underlying these thoughts generalized to conventional resting-state data from the Human Connectome Project (HCP, *N* = 908), revealing a robust mode of population covariation: individuals were predominantly spread along a positive-negative axis linking traits, lifestyles, psychometrics and demographics with a specific pattern of ongoing thoughts predicted from rsFC patterns. This link between behavior and model-predicted thoughts mirrors the pattern previously observed between these behavioral measures and rsFC^15^. Together, these findings suggest that rsFC relates to behavior may in part because it reflects stable differences in individuals’ ongoing thoughts. Our study addresses the scientific gap in understanding the temporal dynamics and patterns of ongoing thoughts and their role in functional brain organization and everyday behavior.

## Results

### Ongoing thoughts during resting-state fMRI

Sixty participants performed up to four 10-minute runs of an annotated rest task as part of a two-session fMRI study that also collected movie-watching, continuous performance task, and post-scan memory and narrative engagement data. Each run of the annotated rest task had 8 trials. In each trial, participants rested for 30 seconds, verbally reported their ongoing thoughts during the preceding resting period for 10 seconds, and rated them on 9 dimensions on a 9-point scale. This 30s rest window was selected based on prior evidence indicating that intervals of this duration reliably capture meaningful fluctuations in cognitive states from FC patterns^46,47^. These dimensions captured wakefulness, thoughts about the external environment, future, past, self vs. others, valence, imagery, word, and specificity^48^ (**Fig. 1**). Ratings spanned the full range of the scale (**Suppl. Fig. 1-2**), suggesting that thoughts during rest vary (see **Suppl. Table 1** for the mean and standard deviation of ratings within- and across participants).

We z-normalized these ratings both within participants and dimensions to control for individual differences and between-dimensional differences within individuals in using the slider bar. To characterize these thoughts at rest, we examined whether and how thought dimensions correlate with one another. 24 out of the 36 possible pairs were significantly correlated within individuals (Bonferroni corrected-*p* < .05; **Fig. 2a**), meaning that dimensions of thoughts are interrelated. For instance, valence was negatively associated with thoughts about the external environment, possibly reflecting scanner-related discomfort, and positively associated with thoughts about the future and others, aligning with theories linking psychological distance to positive affect^49^. Additionally, participants tended to think in the form of images when reflecting on the past, others or positive content, but in words when thinking about the future. Extending prior across-subject findings^37^, we show that these relationships also emerge within individuals over time, demonstrating that the internal architecture of spontaneous thoughts is preserved at the level of moment-to-moment dynamics.

**Figure 2.**
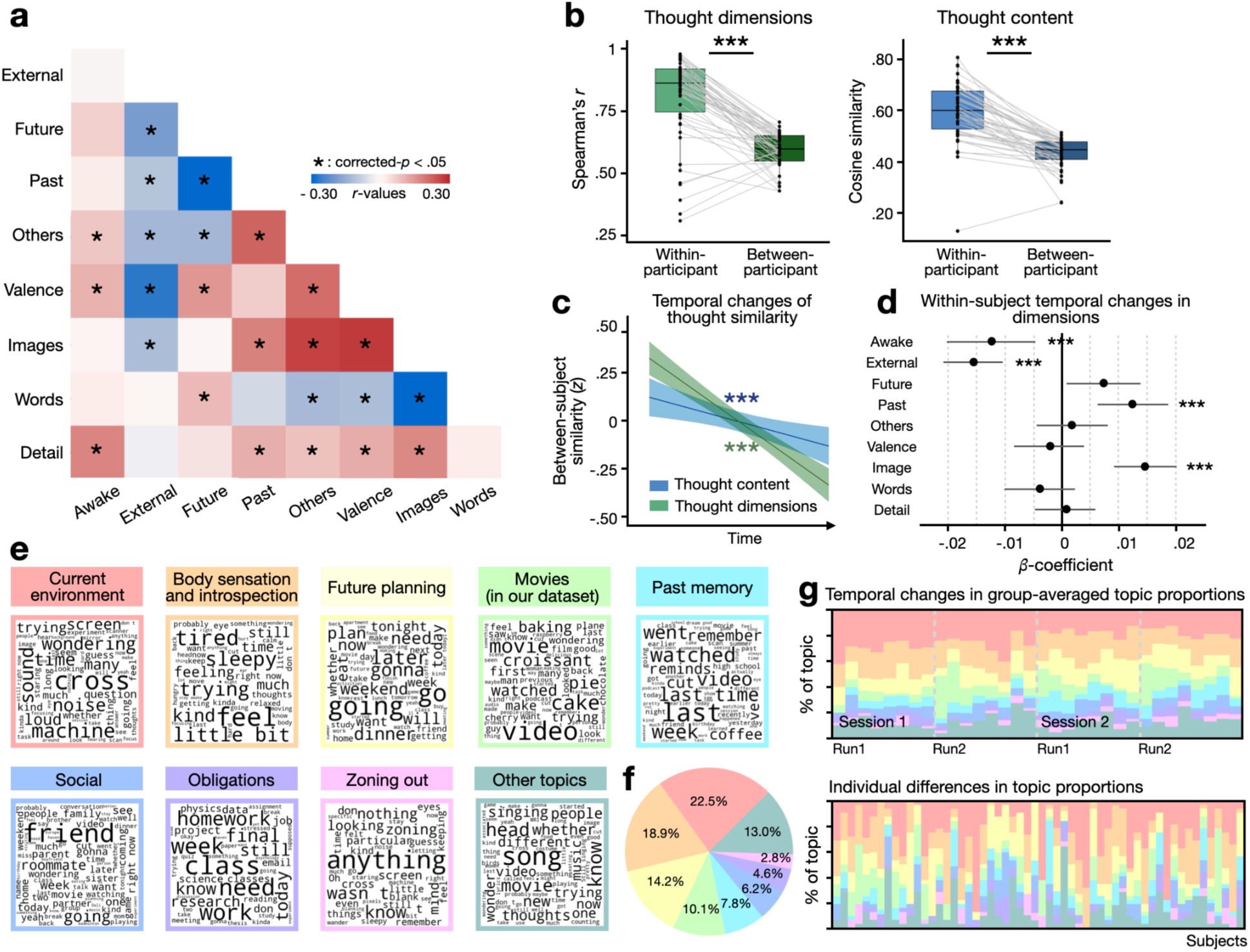
(**a**). Heatmap showing pairwise similarity between thought dimensions. *Bonferroni- corrected *p* < .05 (**b**) Comparison between within- and between-participant thought similarity. Dimensional similarity is measured as Spearman correlation between the 9-dimension self-reported ratings; content similarity measured as cosine similarity between the 512-element sentence embeddings extracted from the speech using Google’s Universal sentence encoder (USE). Dots with connected lines represent individual data points: within-participant (left) and between-participant (right). **(c)** Nested mixed-effects models predicting between-participant similarity (*z*-normalized) from trial number. Negative slopes demonstrate that between-participant similarity in both thought dimensions and content decrease over time. The shaded area represents a 95% confidence interval. (**d**) *Z*-coefficients from nested mixed-effects models predicting ratings on each dimension by trial number (1-32). Positive coefficients indicate increasing ratings, negative indicate decreasing ratings. Violin plots represent null distributions from shuffled ratings (**Bonferroni-corrected *p* < .01). **(e)** Word clouds of frequent terms per thought topic. Larger font indicates higher frequency. **(f)** Proportions of thought topics. Colors match Fig.2e. **(g)** Top: Group-averaged topic proportions change over time, with each column representing a trial. Proportions are calculated as the number of participants thinking about a topic divided by the total number of participants. Dashed lines mark run and session transitions; Bottom: individual-level topic proportions, with each column representing one participant. ***: *p* < 0.001

Next, we tested whether thought patterns are reproducible across scan sessions (mean interval = 10.90 ± 9.61 days). For each participant, we computed the Spearman correlation between their 9-dimensional thought vectors averaged across trials within each scan session and compared the resulting distribution of *r*-values against zero using bootstrapping (10,000 iterations). Thought patterns showed high test-retest reliability (mean *r* = 0.616, 95% CI = [0.526,0.699], *p* < 0.001), indicating that while in-scanner thoughts are dynamic, they are stable across scan sessions and reflect trait-like characteristics.

We further examined whether people show stable individual differences in thoughts above and beyond these group-shared patterns. We compared the similarity of the overall pattern of self-reported thought dimensions between scan sessions from the same individual with those of different individuals. We observed greater within-participant similarity than between-participant similarity, suggesting that individuals have unique thought patterns compared to the averaged others (**Fig. 2b**, left; *t*(53) = 9.96, *p* < .001). Moreover, participants’ thought dimensions became less similar to each other over time (**Fig. 2c**; *β* = -.0205, *s.e.* = .008, *z* = -8.75, *p* < .001), which may reflect attentional shifts from the shared external environment (e.g., scanner noise) towards more spontaneous, internally generated thoughts (e.g., personal memory) as the scan progresses.

The online thought probes enabled us to investigate how dimensions of ongoing thoughts change across trials and scan sessions. For each dimension, we built nested mixed effects models to predict ratings from time (trial number) with intercepts of subjects and sessions nested within subjects included as random effects. Across the 32 trials, people became less awake (*β* = -.013, *s.e.* = .004, *z* = -3.44, corrected-*p* < .01), and thought less about the external environment (*β* = -.016, *s.e.* = .003, *z* = -5.91, corrected-*p* < .01), more about the past (*β* = .012, *s.e.* = .003, *z* = 4.20, corrected-*p* < .01), and more in the form of images (*β* = .014, *s.e.* = .003, *z* = 4.99, corrected-*p* < .01; **Fig. 2d**; see **Suppl. Fig. 3** for within-session changes), revealing shared trends of thought dynamics.

To assess what people were thinking in addition to how they rated it, we analyzed participants’ verbal reports of their thoughts. Semantic meaning of thought content was quantified as 512-dimensional sentence embeddings using Google’s Universal Sentence Encoder (USE). Similarity between text embeddings was assessed with cosine similarity. Replicating the results observed with self-reported thought dimensions, USE-measured thought content showed high test-retest reliability across sessions (mean cosine distance = 0.599, 95% CI = [0.566,0.639], *p* < .001), was more similar within than between participants (**Fig. 2b**, right; *t*(49) = 11.24, *p* < .001) and became more unique over time (**Fig. 2c**; *β* = -.079, *s.e.* = .023, *z* = -3.46, *p* = .001).

To identify thought topics shared across individuals, five independent annotators categorized the free speech from each trial into one of nine possible themes: current environment, body sensation and introspection, movies (that participants watched in the current study), social, future planning, past memory, obligations, zoning out, and other topics (**Fig. 2e**; see **Suppl. Table 2** for instructions to annotators). Inter-rater consistency was high: 94.5% of the thoughts received consensus from at least three annotators. The primary thought topics reflected the current environment, internal feelings and planning for the future (**Fig. 2f**). Echoing the dimensional and content analysis, we observed dynamic shifts in thought topics over time (**Fig. 2g**, top) and substantial individual differences in topic proportions (**Fig. 2g**, bottom; number of topics ranges from 2 to 9, mean = 6.446 ± 1.603).

Together, these findings highlight idiosyncrasy in ongoing thought during resting-state fMRI scans between individuals and over time, both in terms of self-reported ratings as well as the content and topics of thoughts. In the subsequent analyses, we asked how these idiosyncratic thoughts are related to rsFC patterns.

### Thought similarity tracks whole-brain resting-state functional connectivity similarity

Individual variability in whole-brain rsFC patterns has been shown to be robust and reliable, serving as “fingerprints” that can identify individuals and predict their behaviors^5^. First, we examined whether rsFC reflects ongoing thoughts by testing whether individuals with more similar ongoing thoughts have more similar whole-brain rsFC patterns (**Fig. 3a**). Functional connectivity was assessed using a functional brain atlas consisting of 268 regions of interest (ROIs) covering the entire brain^50^. We extracted the ROI-wise BOLD time series from a 30-second window of rest after shifting by 5 seconds to account for the hemodynamic lag and computed the Fisher-z transformed Pearson correlation between the time series of all pairs of ROIs. Functional connectivity similarity was measured as the Pearson correlation between the lower triangles of these 268×268 functional connectivity matrices.

**Figure 3.**
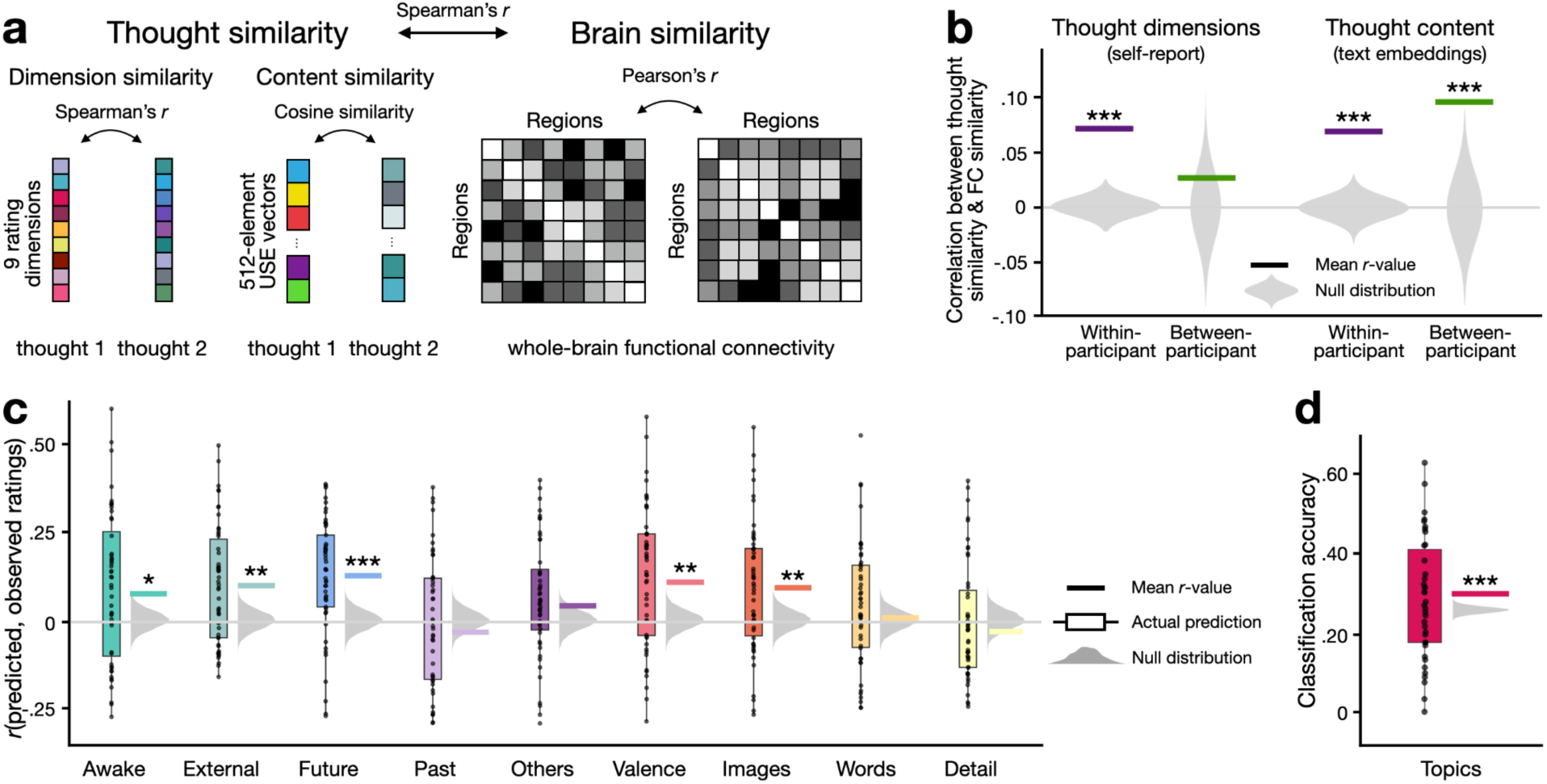
Functional connectivity reflects resting-state thoughts. (**a**) Schematic for representational similarity analysis (RSA) that links thought similarity with brain similarity. Brain similarity was calculated as the Pearson correlation between whole-brain functional connectivity patterns. Thought similarity includes (i) content similarity, measured as the cosine similarity between the 512-element text embeddings extracted using Google USE from the speech describing thoughts from two trials, and (ii) dimension similarity, computed as the Spearman correlation between the 9-dimensional thought ratings from two trials. (**b**) Correlation between thought similarity and functional connectivity similarity. The horizontal lines indicate the actual correlation. The violin plots show the null distributions, generated by shuffling the labels of individuals for the between-individual analysis or the labels of trials within individuals for the within-individual analysis for 10,000 iterations. Statistical significance was assessed by comparing the actual correlation value with the null distribution. (**c**) Functional connectivity predicts dimensions of ongoing thoughts. CPM performance in predicting the 9 thought dimensions. The y-axis represents the predictive accuracy, as measured by Pearson’s correlation between the model predicted ratings and participants’ self-reported ratings. Each datapoint in the box plot represents the predictive accuracy in each round of cross-validation. The gray half-violin plots show the null distribution of 10,000 permutations, generated by shuffling the self-reported ratings before testing the models. Significance was assessed by comparing the empirical mean accuracy against the null distribution. (**d**) Functional connectivity predicts the eight topics of ongoing thoughts. The y-axis represents classification accuracy of the SVC model. Each dot on the box plot indicates classification accuracy in each round of cross-validation, which was measured as the number of correctly predicted trials divided by the overall number of trials. The red horizontal line shows the averaged classification accuracy across all rounds of cross-validation. The null distribution was shown in the gray half-violin plot, generated by shuffling the topic labels within-subjects before testing the model (10,000 iterations). We note that this null distribution is higher than the expected chance level (i.e., 1/8) due to imbalance among thought topics. Please see Suppl. Fig. 9 for balanced accuracy. * *p* < .05, \*\**p* < .01, \*\*\**p* < .001 (Bonferroni-corrected)

Comparing mean functional connectivity matrices between participants revealed that similarity in thought content (*r* = .096, *z* = 4.063, *p* < .001; **Fig. 3b**), but not dimensions (*r* = .026, *z* = .601, *p* = .141), correlated with similarity in rsFC patterns. The *z*-statistic reflects how far the observed correlation deviates from a null distribution generated by permuting thought similarity across participants. In other words, individuals with similar ongoing thought content at rest have similar whole-brain functional connectivity profiles.

Comparing trial-by-trial functional connectivity matrices within participant, we found that trials with more similar thought content (*r* = .069, *z* = 8.919, *p* < .001) and dimensions (*r* = .071, *z* = 10.634, *p* < .001; **Fig. 3b**) had more similar rsFC patterns. This suggests that rsFC dynamics also track within-individual thought fluctuations over time. In addition, this effect extended across subjects: within a given trial, greater between-subject FC similarity tracked greater similarity in concurrent thoughts for both thought content (*r* = 0.0183, *z* = 3.330, *p* < 0.001) and thought dimensions *(r* = 0.0106, *z* = 2.130, *p* = 0.016; **Suppl. Fig. 4**). Thus, inter-individual differences in FC reflect not only stable traits, but also moment-to-moment variations in ongoing experience.

As the annotated rest periods are short and interleaved with thought reports, a critical question is whether FC estimated from these 30-second windows reflects canonical resting-state dynamics. To address this, we compared FC observed during annotated rest with that from conventional resting-state scans and 8 additional task conditions (e.g., working memory, social, language) in the HCP dataset. FC during annotated rest was significantly more similar to HCP rest than to any task condition (all corrected-*p* < 0.001; **Suppl Fig. 5**), indicating that annotated rest preserves connectivity characteristics of conventional rest. Together, these findings reveal a significant link between whole-brain functional connectivity patterns and ongoing thoughts at rest, and suggest that this coupling reflects intrinsic resting-state cognition rather than arising solely from task-driven or demand-related processes.

### Resting-state functional connectivity predicts dimensions and topics of ongoing thoughts

Having established a link between ongoing thoughts and whole-brain functional connectivity patterns, we proceeded to examine whether functional connectivity patterns predict thoughts. We built connectome-based predictive models (CPMs)^51^ with leave-one-subject-out cross-validation (LOSO-CV). In each round of cross-validation, we selected a set of functional connections (FCs) that were significantly correlated with the behavioral ratings of *N*-1 participants in the training set (one-sample *t*-test of within-subject correlations between ratings and FC strength, *p* < .05). We trained a non-linear support vector regression (SVR)^52,53^ model to predict behavioral ratings from the selected features in the training set, and applied the model to FC data from each trial of the held-out participant. Predictive power of each round of cross-validation was assessed as the within-subject Pearson correlation between model-predicted and ground-truth subjective ratings, *R*^2^, and mean squared error (MSE). 50 participants with >20 usable trials (out of 32; each with ≥ 12 TRs post-motion censoring and ≤ 3 missing nodes) were included in the analysis. Despite the relatively brief rest periods and stringent motion censoring, an average of 24.74 TRs per trial remained for estimating rsFC (**Suppl. Fig. 6; Suppl. Table 4**).

We observed significant predictions in 5 out of the 9 thought dimensions (Bonferroni-corrected *p* < .05 for all three evaluation matrices–*r*-values, MSE, and *R^2^*): awake/alert, thinking about the external environment, thinking about the future, valence, and thinking in the form of images (**Fig. 3c**; See **Suppl. Table 5** for statistics). None of these models achieved higher accuracy in predicting any other dimension than itself, indicating model specificity (**Suppl. Fig. 7**). We visualized the FCs that positively and negatively predicted these thought dimensions in **Fig. 4a**. To assess if connections within/between certain functional networks are represented in the neural correlates of thoughts more frequently than predicted by chance, we quantified the proportion of significant FCs within each pair of functional networks and compared them to a null distribution generated by randomly redistributing the same number of FCs across the brain. **Fig. 4b** shows functional network pairs that underlie each thought dimension. For instance, coupling between the default mode and visual networks predicted imagery^54–57^. Together, these results suggest that thought dimensions are encoded in the interactions within and between multiple large-scale functional networks distributed across the brain.

**Figure 4.**
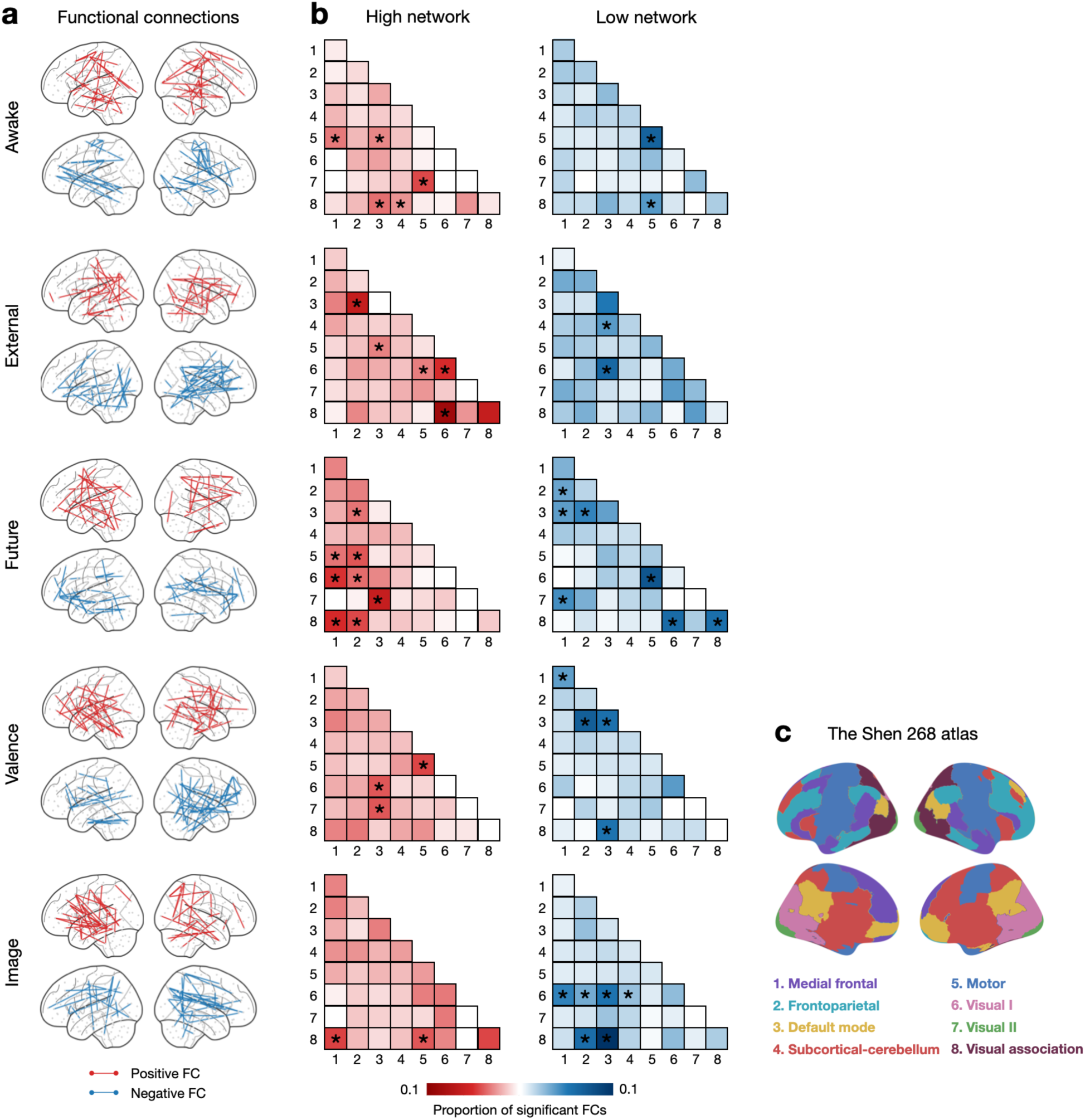
Functional anatomy of thought dimensions with above-chance predictive accuracy. (**a**) FCs that predicted each thought dimension. Red and blue lines respectively represent FCs that positively and negatively predict the corresponding thought dimension. (**b**) The networks that positively and negatively predict each thought dimension. Each cell represents the proportion of selected FCs among all possible FCs between each pair of large-scale canonical functional networks^5^. *: Bonferroni-corrected *p* < .05. **(c)** Visualization of the Shen 268-node atlas. Figure adapted from Finn et al., 2015.

To assess model robustness, we tested whether these models of thought dimensions generalized to predict complementary measures that do not rely on self-report. Indeed, the model trained to predict self-reported wakefulness generalized to trial-to-trial pupil size during rest periods (mean *r* = .128, *p* < .001). The same model also predicted trial-to-trial strength of a validated FC network that positively predicts sustained attention^45,46^ (mean *r* = .049, *p* = .033) in the held-out individual. The model trained to predict self-reported thought valence generalized to predict the probability of negative (mean *r* = -.065, *p* = .010) though not positive (mean *r* = .013, *p* = .313) sentiment of the held-out individual’s free speech extracted from the roBERTa-base model^58^.

We had access to a unique dataset that allowed us to test the out-of-sample generalizability of the imagery model. This sample included fMRI data collected by Megla and colleagues^59^ as an aphantasic individual and their identical twin watched a 10-min non-narrative video *Inscapes*^60^. We hypothesized that our model would predict that the aphantasic individual, who lacks visual imagery, has fewer thoughts in the form of images than their non-aphantasic twin. To test this, we predicted to what extent the thoughts are in the form of images by applying our imagery model to each twin’s rsFC pattern averaged across the movie. As expected, the predicted imagery score on a *z*-scored, standardized scale was negative in the aphantasic twin (-0.31) but positive (0.24) in their healthy twin. Importantly, their difference was significantly above chance as tested with a conservative permutation, where we redistributed the selected edges of the imagery network across the brain by shuffling the node indices before predicting imagery from the FC patterns (*p* = 0.016; **Suppl. Fig. 8**). These results provide evidence for the generalizability of the imagery model and suggest that the aphantasic individual may think less in the form of images.

Finally, we tested if rsFC patterns can distinguish thought topics. Using a LOSO-CV approach, we trained a nonlinear support vector classification model (SVC) on whole-brain FC patterns from *N* - 1 participants to predict thought topic labels (**Fig. 2e**, “other topics” were excluded as they did not form a coherent topic), and applied it to classify thought topics in each trial of the held-out participant. 38 participants with >20 usable trials (out of 32 trials; each with audible speech, ≥ 12 TRs post-motion censoring, ≤ 3 missing nodes) were included in this analysis. The model exhibited above-chance accuracy in classifying thought topics (mean accuracy = 28.5%, *z* = 3.442, *p* = .0003 assessed with permutation; **Fig. 3d**). We note that elevation of the observed null above expected chance level (1/8) reflects imbalances in topic frequency. To account for this, we additionally evaluated classifier performance with balanced accuracy, defined as the mean recall computed across the 8 topics. Using this metric, 36 out of the 38 participants (94.7%) exhibited above-chance performance (mean accuracy = 20.3%, *p* < 0.001 assessed with bootstrapping; **Suppl. Fig. 9**). Together, these results suggest that resting-state functional connectivity encodes the dimensions and topics of ongoing thoughts.

To further test robustness, we re-estimated rsFC using three complementary controls: additional temporal shifts (5 or 10 TRs), two extra censored TRs to account for spin-history effects^61^, and an alternative DVARS-based censoring criterion^62^. Across all controls, rsFC patterns remained highly similar to the original estimates and produced closely comparable results (**Suppl. Fig. 10–12; Suppl. Table 6**). Motion and data-retention metrics (i.e.,, frame-wise displacement, number of censored TRs and temporal signal-to-noise ratio) were likewise unrelated to thought measures and their model predictions at both the within- and between-subject level (**Suppl. Table 7**). Together, these findings argue against hemodynamic lag, motion, spin-history effects, or data quality as explanations for our results.

### Predicted ongoing thoughts at rest are associated with out-of-scanner behavioral measures of individual differences

To establish robust brain-behavior relationships^63,64^, we sought to test the generalizability of our models in an independent dataset. We further asked whether specific patterns of ongoing thoughts at rest are associated with measures of individual differences in behavior. To this end, we applied our thought models to resting-state data in the Human Connectome Project (HCP; *N* = 908) dataset and examined whether model-predicted thoughts at rest meaningfully relate with out-of-scanner individual difference measures.

The predicted thought patterns for each individual included 9 SVR-predicted thought dimensions and SVC evidence for 7 thought topics. The “other topics” was excluded following the same rationale as described above, and the “movie” topic was excluded as the HCP participants did not watch the same movies as participants in our dataset. We used the same set of 478 behavioral measures as Smith and colleagues^15^, which included cognitive abilities (e.g., working memory, pattern comparison processing speed), personality traits (e.g., the five factor), demographics (e.g., age, education), substance use (e.g., tobacco and alcohol) and other behavior (e.g., rule-breaking behavior; see **Suppl. Table 9** for the complete list). We linked model-predicted thoughts with these behavioral measures using canonical correlation analysis (CCA), which identifies the relationship between two sets of variables from their cross-covariation matrices. Specifically, in our case, this technique finds the transformation matrices (A and B) to transform the thought (T) and behavior matrices (S) respectively to U and V that maximizes the correlation between U and V (**Fig. 5a**). To avoid overfitting, we conducted principal component analysis (PCA) on both thought and behavioral matrices before feeding into CCA (**Suppl. Fig. 13**).

**Fig. 5.**
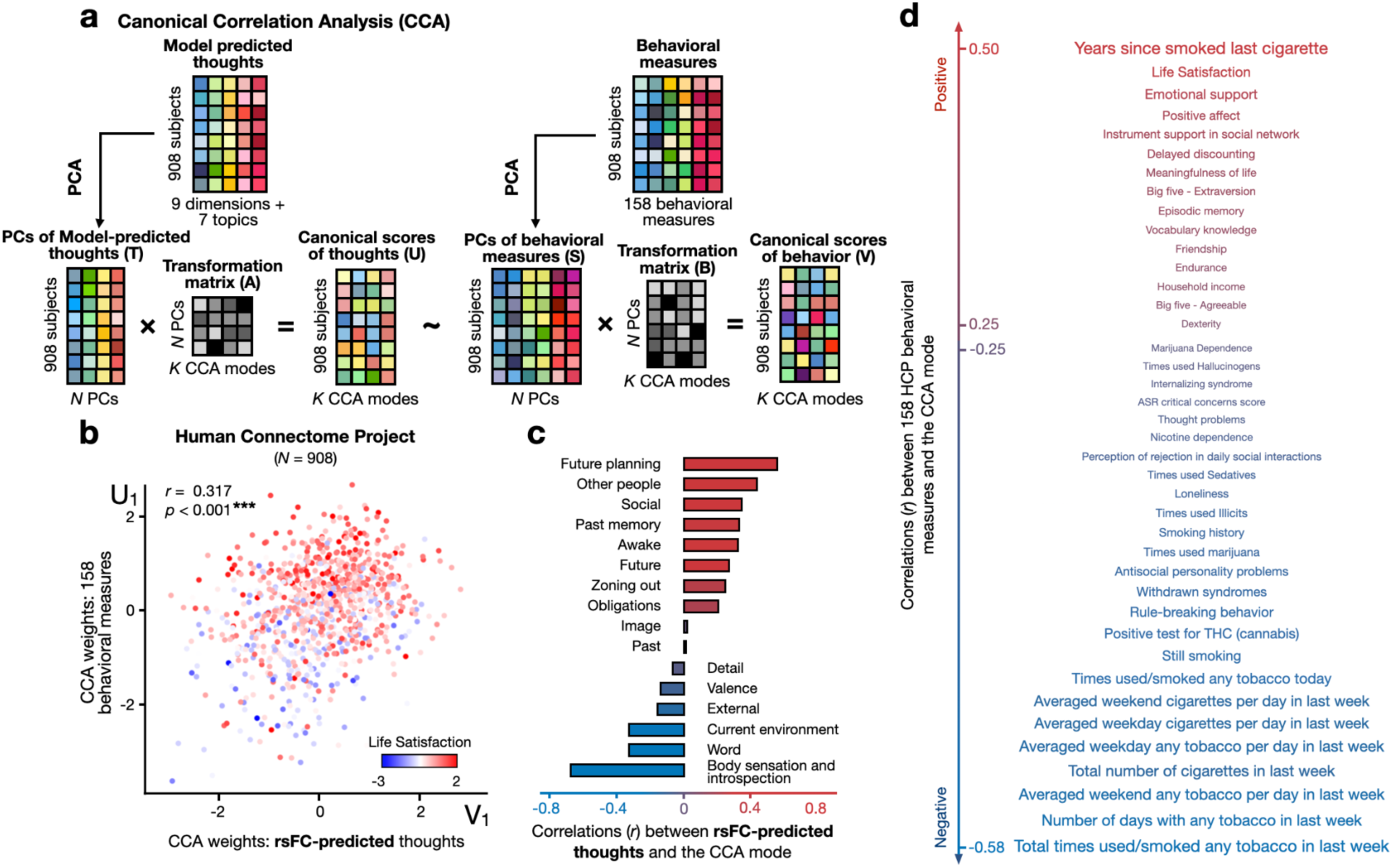
(a) Canonical correlation analysis (CCA) identifies the relationship between two sets of variables — predicted thoughts (T) and behavioral measures (S)—by finding their linear combinations (U and V) that maximize their correlations. **(b)** Correlations between the identified CCA mode and predicted thoughts (x-axis) and behavior (y-axis). Each dot, corresponding to a subject, is colored by a positively associated measure with the CCA mode (e.g., life satisfaction). **: FWE-corrected p < .01. **(c)** Correlations between predicted thoughts and the CCA mode; red and blue represent positive and negative correlations, respectively. Please see **Suppl. Table 10a** for detailed statistics. **(d)** Left: Behavioral measures most strongly associated with the CCA mode in HCP (r > .15). Text color denotes correlation direction; font size reflects explained variance. Please see **Suppl. Table 10b** for the original variable names and detailed statistics.

To successfully link predicted thought patterns with behavioral measures, two prerequisites must hold: First, our thought models must be able to generalize across datasets, capturing meaningful thought differences between individuals from their rsFC patterns. Otherwise, the predictions would merely reflect noise and thus should not meaningfully relate to individual difference measures. Second, ongoing thoughts at rest per se must relate with individual differences^27–36^. If not, the model-predicted thoughts, as a proxy of the actual thoughts, would not relate to individual differences.

We observed one significant CCA mode linking model-predicted ongoing thoughts at rest with individual difference measures (*r* = 0.317, *p* < .001 corrected across all estimated CCA modes; **Fig. 5b**). Significance was assessed by comparing the observed correlation against a null distribution from shuffling subjects while preserving family structure.

Individual difference variables most positively associated with this mode included time since last cigarette, life satisfaction, emotional support, positive affect, and delay discounting. In contrast, variables negatively associated included tobacco use frequency, cannabis (THC) test positivity, rule-breaking behavior, withdrawn syndromes, antisocial problems and loneliness. Together, these form a positive–negative axis, with favorable traits (e.g., good cognitive ability, socioeconomic status, well-being) at one end of the spectrum and adverse variables (e.g., substance use, psychiatric symptoms) at the other (**Fig. 5d**). This pattern, linking model-predicted thoughts with behavior, mirrors prior findings by Smith and colleagues^15^, who linked rsFC directly with the same behavioral measures using the same method (**Suppl. Fig. 14a**). The resulting behavioral weight rankings obtained from our analyses were correlated with those observed in Smith et al., (Spearman’s *r* = 0.757, *p* = 4.58×10^-89^; **Suppl. Fig. 14b**), indicating that these thoughts relate with behavior in a similar manner as how rsFC relates with them. This suggests that the link between rsFC and behavior might reflect differences in ongoing thoughts.

Examining the weights of model-predicted thoughts, we found that future planning, social thoughts, past memory and staying awake showed the strongest positive associations with the identified CCA mode, while body sensation and introspection, thoughts in forms of words and thoughts about the current environment exhibited the strongest negative associations (**Fig. 5c**).

To ensure robustness of our results, we repeated this analysis with varied parameters of dimensionality reduction for both predicted thoughts and behavioral measures (see Methods). In all versions of this analysis, the primary CCA mode shows significantly better fit than the null distribution. In addition and crucially, we investigated whether this effect was specifically driven by edges selected in our ongoing thought models, rather than by any edges. To that end, we built null thought models in our CAT dataset by predicting thought ratings and topics shuffled across trials within participants from rsFC patterns (1,000 iterations; see **Suppl. Fig 15** and **Suppl. Table 11** for the summary characteristics of null-thought models). We estimated CCA fits with predicted thoughts from these null thought models and compared the CCA fit from the real thought model with this conservative, new null distribution. Again, the observed CCA fits remain relatively stable and outperformed the 95% percentile of the null models across multiple combinations of thought and behavior PCs (paired-*t* = 3.393, *p* = 0.002; **Suppl. Fig. 16**).

Finally, although our primary CCA model incorporated predictions from all nine dimensional decoders with the topic decoder, the effect remained robust across alternative rsFC model specifications—including dimensional-only, topic-only, and other multidimensional decoder variants (**Suppl. Table 12**)—indicating that the rsFC–behavior association is not driven by any single set of thought features.

Taken together, these results provide strong evidence for model generalizability and suggest that ongoing thoughts at rest are associated with measures of individual differences in behavior above and beyond the effects of rsFC per se. Ongoing thoughts at rest may therefore contribute to rsFC-behavior associations.

## Discussion

Resting-state fMRI has been widely used in cognitive neuroscience research, playing a crucial role in large-scale neuroimaging initiatives seeking to map functional brain architecture and relate it with individual characteristics. Yet, we still lack a clear understanding of what the brain is doing at rest. In particular, it remains largely unexplored how ongoing mental experiences during rest shape brain network dynamics. To contribute to filling this gap, the present study utilized the annotated rest task, which regularly samples individuals’ ongoing mental experiences during resting-state fMRI. Our findings demonstrate that resting-state functional connectivity patterns indeed reflect rich and individualized ongoing thoughts. Neuromarkers of these thoughts generalized to independent samples, tracking individual differences in cognitive abilities, psychiatric symptoms, personality traits, lifestyles, and demographics. Together, our findings highlight ongoing mental experiences as a notable factor in understanding functional brain organization, everyday behavior, and their associations.

Functional networks have traditionally been thought to reflect stable individual characteristics^5^, with limited attention paid to the influence of spontaneous cognitive fluctuations during rest^65,66^. Our findings challenge this conventional view in the following ways. First, we demonstrate that ongoing thoughts can be reliably decoded from rsFC patterns, suggesting that ongoing brain network dynamics track these in-scanner mental experiences. Second, our findings support the observations that individuals with more similar overall self-reported thoughts at rest exhibit more similar rsFC profiles^37^ and extend these findings by demonstrating that within-individual thought fluctuations also track rsFC dynamics. Third, although ongoing cognition is often viewed as too variable to contribute meaningfully to the relatively stable patterns of rsFC, our results resonate with recent findings that individuals exhibit more stable patterns of in-scanner thoughts across scan sessions than originally assumed^67,68,37^. This high test-retest reliability of resting thoughts helps reconcile how individuals exhibit trait-like thought patterns that relate to rsFC, despite its intrinsically dynamic nature. Thus, these results together provide strong evidence that systematic differences in ongoing mental experiences are reflected in systematic differences in rsFC.

Our work builds on a rich literature demonstrating that diverse aspects of human cognition, such as mental imagery^69–76^, language^77–80^, decision making^81–84^, can be reliably decoded from brain activity. While prior studies have primarily focused on externally driven tasks, our study shows that dimensions and semantic content of ongoing thoughts can be decoded during task-free rest, extending the existing literature by suggesting that the neural representations of cognition are sufficiently robust and intrinsic to be recoverable even when they unfold spontaneously without external control or experimental scaffolding. In addition, the success of our connectome-based approach complements prior literature on brain activation patterns and contributes to the growing recognition that dynamic interplays between large-scale brain networks support cognition^85–87^.

Our findings are consistent with previous studies demonstrating that patterns of ongoing thoughts are predictable from rsFC^36–38^. For example, a recent work by Gonzalez-Castillo and colleagues showed that individuals’ retrospectively reported experiences—such as wakefulness, imagery, valence, and surrounding focus—can be decoded from rsFC. Our results replicated and extended this prior observation, indicating that not only broad self-rated dimensions but also the specific semantic meanings of ongoing thoughts can be reliably predicted from rsFC, both across individuals (trait-level differences) and within individuals over time (state-level fluctuations). The content of thoughts—rich in personal goals, concerns, memories, and emotions^3,26–31^—offers a window into internal states and are predictive of cognitive traits and psychiatric symptoms^32–35^. However, assessing them remains challenging: they arise freely, often without conscious control, and the act of observing them can alter their nature. Inferring their content directly from brain activity thus offers a promising alternative for accessing internal experiences without disturbance.

A key strength of our work lies in the generalizability of our thought-predictive models, which capture the patterns and content of individual spontaneous thinking from these identified neural substrates. Our models, generalized to non-introspective measures (e.g., pupil size, free speech sentiment) and out-of-sample individuals, offer a foundation for broader application. For example, probing thoughts is challenging in healthy individuals, and even more so in clinical populations. Our ability to identify aphantasia using the imagery model suggests that future work may similarly detect impaired prospection or depressive affect using our models of future thinking and valence. Moreover, in response to recent calls for more contextualized and behaviorally meaningful frameworks beyond the traditional resting-state paradigms^43^, incorporating model-predicted in-scanner thoughts may improve interpretability of nebulous in-scanner experiences, for example, during post-social rest^88,89^, mood drifts^90,91^, and narrative comprehension^92,93^. We will publicly release the trained model weights and analysis code (see Data and Code Availability), such that other researchers can apply the models to their own data or publicly available fMRI datasets.

In addition, our findings offer insights into biomarkers of spontaneous thinking during rest. We demonstrate that such thinking recruits brain regions involved in high-level reasoning, sensory processing and memory^17–19,94,95^. For instance, we observed increased interactions between the medial frontal, default model, frontoparietal and visual network as people think about the future, consistent with prior studies linking these systems to future thinking^96–99^. Similarly, coupling between the default mode and visual network tracks thinking in the form of images, echoing earlier findings on the neural correlates of imagery^54–57^. Notably, the default mode network—a set of brain regions consistently more active at rest than during experimental tasks^95,100^—is heavily involved in all observed thought dimensions, reinforcing the perspective that it supports the generative dynamics of mind-wandering rather than merely reflecting a task-free state^2,4,101,102^.

We extended neuroscientific investigations of thought-related rsFC dynamics to its individual differences. To our knowledge, these findings are the first to demonstrate that neural signatures of ongoing thoughts are associated with individual differences in behavior. Building on prior work^15^ linking rsFC with these measures, our results suggest that such associations are partly driven by individual patterns of spontaneous thoughts at rest. While prevailing theories attribute the rsFC-behavior associations to intrinsic differences in the efficiency of brain connectome architectures—independent of any ongoing cognition—that support behavior (e.g., attentional performance^45,46^), we offer a complementary hypothesis: rsFC predicts individual differences in behavior may be partly because people with different behavioral tendencies engage in different patterns of thinking during rest. Our findings echo a recent growing cognition^37^ that underscores a crucial consideration for studies mapping traits or clinical diagnoses to brain connectomes: interpretations of rsFC must account not only for intrinsic network architecture but also for the brain’s spontaneous activity during rest. Incorporating individuals’ in-scanner mental experiences may thus practically enhance the predictive power of rsFC for behavior.

We point out a few limitations of our work

First, the optimal frequency and format of thought probes remains an open question^16,43^. In our current experiment, participants briefly described their thoughts aloud after each 30s rest period, followed by ratings on nine dimensions. Compared to prior approaches that rely on binary mind-wandering probes (e.g., on/off-task)^2,102^ or dimensional ratings^22,27,36,37^, this design offers a richer and more fine-grained view of these thoughts and thus allows for more granular mapping between the fluctuating mental^16^ and brain states^102,103^. However, a key limitation of frequent probing is that it can affect the flow of thoughts, which raises the possibility that the observed thoughts and their corresponding brain dynamics may differ in their nature from longer, undisrupted rest. Several findings mitigate this concern. First, the reported thoughts were highly variable across individuals and time, suggesting that they retain their spontaneous, idiosyncratic nature. The content of these thoughts echoes earlier findings based on meta-analytic decoding of rsFC^104^, which similarly identified recurrent themes such as episodic memory, planning, social cognition, inner speech, somatosensory processes, and visual imagery as dominant components of spontaneous cognition during rest. Second, the FC patterns observed during the 30-second rest period in the annotated rest task are more similar to HCP rest than any other tasks. Third, models trained on these data related to individual difference measures in the HCP dataset—where no thought probes were present—suggesting that the observed associations between rsFC and behaviorally relevant mental states are not just an artifact of the probing process. Nonetheless, we cannot exclude the possibility that participants habituate to predictable thought probes, such that anticipatory processes shape ongoing mentation. Prior studies have used longer inter-probe intervals (e.g., 45–90s) and randomizing probe timing^24,38,105–108^, and modeling probe-related variance through nuisance regression^20^. However, these studies did not examine how such design features shape the relationship between ongoing thoughts and brain dynamics. Future work should therefore systematically test how probe presence, frequency, timing, and format influence reported thoughts and their coupling to brain dynamics.

Second, although our thought models achieved statistically significant predictive power, their accuracy was relatively modest. Yet, the theoretical value of a robust and reliable model does not hinge on predictive precision alone^63^. Importantly, when applied to HCP dataset, model-predicted thoughts exhibited structured associations with independent behavioral measures that were absent in shuffled thought models, suggesting that these models capture domain-general variance in ongoing thoughts that transcends individuals and scanning sites. However, a key limitation is that the HCP dataset does not include in-scanner thought reports, which precludes item-level validation of specific predicted thought dimensions. Directly testing whether predicted dimensions correspond to actual ongoing thoughts in independent samples with concurrent experience sampling is an important direction for future work. In addition, the small effect size cautions against interpreting these model-predicted thoughts as precise proxies for individuals’ in-scanner experiences in new samples. Furthermore, while our models predicted thought fluctuations in held-out participants within the same dataset, their capacity to track moment-to-moment variations across datasets remains untested. Future work is needed to evaluate their temporal precision in new contexts.

Third, our findings suggest ongoing thoughts at rest modestly contribute to the rsFC–behavior association, but we do not claim these thoughts are the only or primary driver of rsFC patterns and its relevance with behavior^37^; other neurobiological and cognitive processes—such as homeostasis^65^, interoception^103^, learning and memory^104,109,110^—might explain more variance. A comprehensive account requires future work to disentangle the relative contributions of these factors. In addition, one-dimensional physiological signals like arousal may provide a global scaffold for large-scale spatiotemporal brain dynamics^111^, raising the possibility that it also contributes to these reported effects. Future studies should directly test this possibility by measuring ongoing thought and concurrent physiological arousal during resting-state fMRI. Finally, many of our analyses are correlational, leaving the inference or interpretation of the causal relationships between thoughts, rsFC patterns and behavior unresolved. For instance, do certain thought patterns causally shape functional architectures, or do intrinsic connectome properties constrain and enable particular ways of thinking? Future studies could examine whether rsFC mediates the link between ongoing thought at rest and everyday behavior.

Despite these caveats, the current study underscores a critical yet often overlooked consideration for the large-scale resting-state fMRI initiatives in cognitive neuroscience research. Our findings provide strong evidence that the mind’s rich ongoing thoughts, as a unique lens into individuals’ internal mental world, are reflected in rsFC patterns and linked to human everyday cognition and experiences. Our work lays the foundations for future investigations into the neural substrates of spontaneous thinking and its critical role in bridging individual differences in functional brain organization and behavior.

## Methods

### Subjects

Sixty healthy individuals from the University of Chicago and surrounding community (34 women, 26 men, mean age 22.86 ± 3.27 years) participated in the study. All participants were right-handed, spoke fluent English, and reported normal hearing and normal vision or corrected-to-normal vision. Participants provided informed written consent before the start of the study in accordance with the experimental procedures approved by the Institutional Review Board of the University of Chicago and were compensated for participation with cash ($130) or course credits (3 credits + $100).

### Experimental procedures

The experiment consisted of two 3-hour experiment sessions. Participants completed the sessions on two separate days, with an average time interval of 10.90 (± 9.61) days between sessions. Four participants did not come back for the second session after completing the first. Each session consisted of an in-scanner phase and a post-scan behavioral experiment phase. Each in-scanner phase included two annotated rest, two naturalistic tasks (movie watching and/or story listening), and an audio-visual continuous performance task (avCPT)^112^ run. The narratives include an audio-only podcast about baking pies (aNT, 16min 33s), a visual-only video about baking croissants (vNT, 16 min 58s), an audio-visual video about baking a cake (avCake, 15min 40s), and a segment of an audio-visual American spy thriller, North by Northwest, directed by Afred Hitchcock (avNorth, 14min 49s). Participants completed the tasks in the following order: single-modality naturalistic task (aNT or vNT), the first annotated rest, audiovisual movie watching (avCake or avNorth), the second annotated rest, and avCPT. The order of aNT and vNT across sessions as well as the order of avNorth and avCake across sessions were counterbalanced across participants. The avCPT and movie data were not analyzed in this study.

Each run of the annotated rest task lasted for 10 minutes. In each run, participants completed eight trials in which they rested for 30 seconds, verbally reported their ongoing thoughts for 10 seconds, and then rated their thoughts on nine dimensions^48^ using a slider bar on a 1-9 scale. The instruction for the verbal report was “Briefly describe your thoughts just before this. Please speak loudly!”. Participants used a response box to rate. The ratings started at 5 at the beginning of each trial and participants moved the slider rightward (up to 9) by pressing with middle finger and leftward (down to 1) by pressing with index finger. The full list of the 9 rating questions and corresponding labels of the right and left extremes of the scale is as follows:

Q1: How awake/alert do you feel? (Not at all awake/alert – Completely awake/alert)

Q2: My thoughts were related to the external environment around me. (Not at all – Completely)

Q3: My thoughts were about the future. (Not at all – Completely)

Q4: My thoughts were about the past. (Not at all – Completely)

Q5: My thoughts were… (About myself – About other people)

Q6: My thoughts were… (Very negative – Very positive)

Q7: My thoughts were in the form of images. (Not at all – Completely)

Q8: My thoughts were in the form of words. (Not at all – Completely)

Q9: My thoughts were detailed and specific. (Not at all – Completely)

The behavioral ratings were z-normalized within participants and dimensions to control for the subjectivity in ratings on the scales. For each dimension, the participants’ ratings were widely distributed (**Suppl. Fig. 1-2**), suggesting a richness in thoughts during rest.

Participants practiced the annotated rest task with an experimenter before scanning to become familiar with the 10-second speaking duration, rating questions, and slider-bar manipulation.

A 90-min post-scan behavioral assessment immediately followed the in-scanner tasks in each session in a separate behavioral testing room near the scanner. This behavioral session consisted of two phases. In the first phase, participants completed a self-paced recognition memory task for the stimuli presented in avCPT^113^. In the second phase, participants separately rated their overall engagement, answered memory questions, and performed a continuous engagement rating task for the single-modality naturalistic stimulus (aNT or vNT) and the audio-visual movie (avNorth or avCake) that they just watched and/or listened to in the scanner. These behavioral data are not included in analysis for this paper.

### Measures of ongoing thoughts during rest

We measured resting-state ongoing thoughts in three ways: dimensions, content, and topics.

#### Thought dimensions

In each trial of the annotated rest task, participants rated their thoughts during the resting period on the 9 dimensions as described above: awake/alert, external, future, past, others, valence, image, word, as well as detailed and specific.

#### Thought content

Three annotators independently listened to the recordings of the speech data and transcribed the audio to text. These transcriptions were encoded as 512-dimensional vectors using Google’s Universal Sentence Encoder (USE), a natural language processing technique implemented in TensorFlow that captures the semantic meaning of the text^114^ (https://www.tensorflow.org/hub/tutorials/semantic_similarity_with_tf_hub_universal_encoder).

#### Thought topics

Five annotators independently read through each of the transcribed thoughts and labeled the thoughts as one of nine pre-determined topics that best fit. To generate these nine topics, we referenced past literature^16^ and derived them from listening to participants’ verbal responses. The nine topics were: current environment, body sensation and introspections, movies, social, future thinking, past memory, obligations, zoning out, and other topics. See **Suppl. Table 2** for detailed definitions, which was the instructions for the annotators to label the thoughts, and **Fig. 2f** for behavioral characteristics (i.e., word clouds) of these topics. At least three annotators agreed on the topic for 94.50% of the thoughts (5 agreeing: 39.7%, 4 agreeing: 28.8%, 3 agreeing: 26.2%). Agreement from at least 3 annotators (out of 5) would suggest consensus from the majority. Thus, for each of these thoughts, the topic with the most votes was set as the ground-truth topic. A small proportion, 5.50%, of the thoughts were agreed upon by only two annotators. Author J.K. went through each of these thoughts and decided on the topic himself. No thoughts had five different opinions from the five annotators.

### Nested mixed effect models on thought dimensions and between-participant thought similarity

To investigate how ongoing thoughts change over time across individuals, we built nested mixed effects models to predict the thought dimensions from trial number (from 1 to 32), with intercepts of subjects and sessions nested within subjects included as random effects. We built separate models for each thought dimension and corrected for multiple comparisons. The models were implemented in python as ‘mixedlm’ with statsmodels.api (https://www.statsmodels.org/stable/api.html). A positive ***β*** coefficient of the variable trial number indicates the ratings increased over time, whereas a negative ***β*** coefficient indicates that the ratings decreased over time. We corrected for multiple comparisons using Bonferroni correction (https://www.statsmodels.org/dev/generated/statsmodels.stats.multitest.multipletests.html).

To examine whether individuals’ thought patterns become more dissimilar to each other across trials and scan sessions, we built nested mixed effects models to predict between-participant thought similarity from trial number (from 1 to 32), again with subject ID and session number nested within the subject as the random variables. Two models were built, one for similarity in thought dimensions and the other for similarity in thought content. The similarity in thought dimensions was measured as the Spearman correlations between the two 9-dimensional ratings. The similarity in thought content was measured as the cosine similarity (1 - cosine distance) between text embedding vectors. For each participant, on the trial level, we calculated their thought similarity with every other participant and computed the group averaged similarity score (Fisher-z transformed for thought dimensions). A positive ***β*** coefficient means that individuals’ thought patterns become more similar to each other over time, whereas a negative ***β*** coefficient means that individuals’ thought patterns become more unique over time.

To examine how thought patterns change across trials within sessions, we repeated this analysis, predicting between-participant thought similarity from trial number (1 to 16) with the intercept of subject ID as the random variable (**Suppl. Fig. 3**).

### Comparing within- and between-participant thought similarity

To examine individual differences in ongoing thought, we compared within-participant and between-participant thought similarity. For thought dimensions, we averaged the 9-dimensional vectors across trials within sessions and participants. For thought content, we combined the transcriptions of free-speech from all trials within session and participant before extracting the 512-dimensional vector of text embeddings. Within-participant thought similarity was measured as the similarity of thought patterns between session 1 and session 2 of the same participant (Spearman correlation for thought dimensions and cosine similarity for thought content). To calculate between-participant thought similarity, for each participant we computed the similarity in thought patterns between one session (i.e., session 1) of this participant and the other session (i.e., session 2) of every other (*N* - 1) participant. The similarity scores were averaged across the *N* - 1 participants. A paired *t*-test was conducted to compare within- and between- participant thought similarity.

To examine whether individuals have the most similar thoughts to themselves as compared to any other participants, when calculating between-participant thought similarity, we took the highest similarity score in the group of *N* - 1 participants rather than taking their mean score. We again conducted a paired *t*-test to compare within-participant similarity with the highest between-participant thought similarity.

### MRI acquisition and preprocessing

MRI data were collected on a 3T Philips Ingenia scanner at the MRI Research Center at the University of Chicago. Functional images were collected with a 32-channel head coil (TR/TE = 1,000/28 ms, flip angle 62°, whole-brain coverage 27 slices of 3 mm thickness, in-plane resolution 2.526 × 2.526 mm^2^, matrix size 80 by 80, FOV 202 × 202 mm^2^, and MultiBand SENSE factor 3). Structural images were acquired using a high-resolution T1-weighted MPRAGE sequence (1 mm^3^ resolution).

We preprocessed the fMRI data using Analysis of Functional Neuroimages (AFNI version 19.0). The first 3 TRs of each functional run were excluded from analysis. Data were then despiked, slice-time corrected and motion-corrected as described previously^115^. Covariates of no interest were regressed from the data, including a 24-parameter head motion model (6 motion parameters, 6 temporal derivatives, and their squares) and mean signal from subject-specific eroded white matter and ventricle masks and the whole brain. Extra motion censoring was performed for annotated rest, where volumes were censored if they contained outliers more than 10% of the voxels or the Euclidean distance of the head motion parameter derivatives were greater than .25mm. We then registered the functional images to participants’ skull-stripped MPRAGE anatomical images with linear transformation and then normalized to the Montreal Neurological Institute (MNI) space with nonlinear warping.

### Functional parcellation and computing functional connectivity

We parcellated the brain using the 268-ROI Shen atlas^50^ and averaged the blood-oxygen-level dependent (BOLD) time courses of all voxels in each ROI. We extracted BOLD time series from a 30-second window starting from the 5th TR of the rest period to account for the hemodynamic lag. We computed the Fisher-z transformed Pearson correlation between the time series of all pairs of ROIs when there were no fewer than 12 available TRs after motion censoring in the 30-second window. Trials that did not meet this criteria were excluded from further analysis due to excessive motion. A 268×268 functional connectivity matrix was calculated for each resting period (trial) for each participant.

### Comparing FC patterns between CAT annotated rest and HCP tasks

To establish that brain activity observed during the brief 30-second rest periods in the annotated rest task is comparable to that during longer, uninterrupted resting-state scans, we compared the FC patterns between CAT annotated rest and rest and 8 additional tasks in HCP. The HCP tasks include working memory, relational, motor, social, emotion, gambling, language tasks, and movies. For each CAT subject, we averaged the FC matrices across trials to estimate an static FC profile. We acquired the HCP preprocessed brain data of all tasks from Song et al.^116^. For each HCP subject (*N* = 119), we compared the FC matrix of each task with the rest FC matrix of every CAT subject using Pearson correlation and averaged across all CAT subjects with Fisher-*z* transformation. We conducted paired-*t* tests between rest-task pairs and corrected for multiple comparisons across all possible pairs.

### Correlating thought similarity with whole-brain functional connectivity similarity

To establish an association between ongoing thoughts and resting-state whole-brain functional connectivity patterns, we first tested whether thought similarity was correlated with whole-brain functional connectivity similarity by conducting a representational similarity analysis (RSA). We performed the analysis on two levels:

1. “between-participant”: on this level, we asked whether static functional connectivity profiles reflect individuals’ overall patterns and content of ongoing thoughts at rest. For each participant, we averaged the functional connectivity patterns across up to 32 rest periods from 4 fMRI runs, combined the free-speech data across trials before extracting text embeddings (for thought content) and averaged the ratings on the 9 dimensions across trials (for thought dimensions). We calculated between-participant similarity in both thoughts and functional connectivity, and computed a Spearman correlation between the two similarity matrices (**Fig. 3a**). The statistical significance of this *r*-value was assessed by comparing it to a null distribution, generated by shuffling the labels of participants in the thought similarity matrix before correlating with the functional connectivity similarity matrix (10,000 iterations). Note that in this particular analysis, we used raw dimensional ratings rather than the z-scored ratings within subjects and dimensions, following the rationale that the mean of z-scored ratings will center around zero. Confounds like rating biases may be inherently embedded in this thought measurement, which may contribute to the nonsignificant results when relating similarity in the mean rating dimensions with similarity in FC profiles. All other analyses involving the dimensional ratings were conducted on z-scored values.
2. “within-participant”: on this level, we asked whether functional connectivity dynamics track within-participants thought fluctuations over time. For each participant, we calculated similarity for thoughts (i.e., thought dimensions and thought content) and functional connectivity patterns between rest periods of each trial and computed the Fisher-z transformed averaged *r*-value. These *r*-values were averaged across participants and compared to a null distribution for statistical significance tests. To generate this null distribution, we shuffled the labels of trials in the thought similarity matrix within participants, correlated it with the functional connectivity similarity matrix, and averaged the null *r*-values across participants (10,000 iterations). We conducted this analysis on similarity in both thought dimensions and content.

### Connectome-based predictive modeling of thought dimensions

We built connectome-based predictive models (CPMs) to predict the dimensions of ongoing thoughts from whole-brain functional connectivity (FC) observed during the intermittent rest periods. Trials with fewer than 12 available TRs after motion censoring and/or more than 3 missing nodes were excluded. We note that the 3 missing nodes represent partial brain cutoff during the scan, usually due to limited slice coverage or slice positioning and most commonly affecting the recording of cerebellum activity. This instance occurred in 18 annotated rest scans centered in 8 participants. All affected scans were excluded from subsequent analyses. All trials across the four runs of the annotated rest task were concatenated for each participant, and participants with more than 20 available trials were included for further analysis. 50 participants remained in the predictive modeling process, where we built CPMs with a leave-one-subject-out cross-validation approach.

We followed the general analytic approach described in Shen et al.^51^ to build the CPMs. For each round of cross-validation, we performed feature selection by identifying FCs that were significantly correlated with behavioral ratings in a training set of *N* - 1 participants (one-sample *t*-test, *p* < .05). We trained a non-linear support vector regression model (sklearn.svm.SVR, https://scikit-learn.org/stable/modules/generated/sklearn.svm.SVR.html, kernel = “rbf”) on the training set to predict the behavioral ratings from the selected FC features. In contrast to “traditional” CPM which relates mean FC strength to behavior with a linear regression, we elected to use SVR to consider the pattern of FCs related to behavior as in previous work^52,53,119^. The SVR model was then used to predict the behavioral ratings for each trial of the held-out participant, and the predictive accuracy was calculated as the Pearson correlation between the model predicted ratings and participants’ self-reported ratings. Correlation coefficients from all cross-validation rounds were Fisher’s *z*-transformed, averaged across all rounds, and transformed back to an average *r*-value to serve as the final measure of the model performance.

To evaluate statistical significance, model performance was compared against a null distribution generated by testing the model predictions on the shuffled behavioral ratings of each participant before averaging these *r*-values across participants (10,000 iterations). Same as the real models, when evaluation the null model performances we Fisher’s *z*-transformed the correlations between model-predicted ratings and shuffled behavioral ratings, averaged them across all cross-validation rounds, and transformed the mean *z*-value back to the averaged *r*-value. We used a one-tailed significance test, with *p* = (1 + number of null *r* values ≥ empirical *r*) / (1 + number of iterations). Separate CPMs were built for each thought dimension. Thus, the *p*-values were corrected for multiple comparisons across 9 dimensions (Bonferroni correction, https://www.statsmodels.org/dev/generated/statsmodels.stats.multitest.multipletests.html).

### Functional anatomy of the thought-dimension CPMs

We investigated the neural correlates of the thought dimensions. For each thought dimension (e.g., wakefulness), we defined the set of FCs selected in every round of cross-validation during the modeling process as the wakefulness network. The wakefulness network consists of high- and low-wakefulness networks, which represent the set of FCs that positively and negatively predicted wakefulness ratings, respectively.

To understand how FCs within and between these large-scale functional networks constitute the high and low FC networks underlying each thought dimension, we grouped whole-brain FCs in 8 canonical large-scale functional networks predefined by Finn et al.^5^: Medial frontal, frontoparietal, default mode, subcortical-cerebellum, motor, visual I, visual II, and visual association networks, and examined whether particular functional networks were represented in the thought dimension FC networks more frequently than chance. We calculated the proportions of selected FCs relative to the total number of possible connections between each pair of functional networks. These proportions were compared to a null distribution, generated by randomly shuffling the positions of the selected FCs (while keeping the same number) within the entire set of FCs. The statistical significance of each proportion was assessed by comparing the actual value to the null distribution of 10,000 permutations with Bonferroni corrections. The FC networks underlying each thought dimension were plotted in **Fig. 4**, with high and low FC networks respectively plotted in the upper and lower triangles.

### Validating the awake and valence models

As an assessment of model robustness, we examined whether the CPMs trained on individual self-report ratings of wakefulness and thought valence generalized to predict complementary measures that do not rely on self-report. Specifically, for the awake dimension, we tested if the awake CPM predicted trial-to-trial pupil size, an indicator of physiological arousal, and strength of the sustained attention network. For the valence dimension, we tested if the valence CPM predicted trial-to-trial positive and negative sentiment of the speech. We tested the awake and valence (also imagery, described in the next section) dimensions, but did not attempt on the other significant dimensions (i.e., external environment and future) as their validations were not easily accessible in the current study setting.

The pupil data was recorded using an Eyelink 1000 eye tracker (SR research) at a sampling rate of 500 Hz. We calibrated and validated the eye tracker with a 9-point calibration at the beginning of each fMRI task, including the annotated rest tasks. We performed linear interpolation on the data, spanning the period from 150 ms before the onset of each blink or saccade to 150 ms after the end of the blink. We smoothed the interpolated data with a zero-phase low-pass filter (third-order Butterworth, cutoff = 4 Hz) and *z*-normalized it within each participant^120^. The pupil diameter time course during the 30s resting period in each trial was averaged as a proxy of physiological arousal in that trial. Pupil diameter was significantly correlated with self-reported wakefulness (*r* = .094, *p* < .001).

To validate the wakefulness model on pupil size data, we trained the awake CPM in the same leave-one-subject-out manner as described above. The only difference was that in each round of cross-validation we tested the model predictions in the held-out participant on their trial-to-trial mean pupil size instead of the self-reported awake ratings. The validation performance was assessed as the averaged predictive accuracies across all cross-validation rounds, which was compared to a null distribution generated by randomly shuffling trial-wise pupil size values within each participant before testing the model and averaging the predictive accuracies. We repeated the same analysis pipeline for testing the wakefulness CPM on trial-to-trial strength of the high-sustained attention network (described below) and for testing the valence CPM of positive and negative sentiment.

The high-sustained attention network was acquired from Rosenberg et al.^45,46^. These studies built connectome-based predictive models to predict individuals’ performances on sustained attention tasks and showed that the identified neuromarkers reliably predicted changes in sustained attention across individuals, datasets, and at various timescales. The high-sustained attention network consists of 757 FCs that positively predict sustained attention. We extracted the FC patterns from the 30s resting period in each trial, and calculated the mean strength of the high-sustained attention network as the averaged strength of the 757 FCs in this network.

We tested the valence CPM on the linguistic sentiment of participants’ free speech. Positive and negative sentiment was extracted from speech data using a pre-trained Twitter-roBERTa-based model from the Hugging Face library (https://huggingface.co/cardiffnlp/twitter-roberta-base-sentiment). This particular model was chosen because it was trained on 58M tweets, which were of similar length and data type as the self-reported thoughts in the current annotated rest task. The model outputs the probability of each thought to be positive, neutral and negative. We applied the model to the written transcriptions of the speech in each trial. Both positive (*r* = 0.302, *p* < .001) and negative sentiment (*r* = -0.440, *p* < .001) are significantly correlated with subjective ratings of valence.

### Validating the imagery model in an aphantasic twin dataset

As an external validation of the imagery model, we examined whether an aphantasic individual, who lacks visual imagery, shows weaker imagery network strength relative to their identical non-aphantasic twin. The functional and structural MRI data were collected by Megla and colleagues^59^ and preprocessed using the same AFNI pipeline as described above.

Each of the two individuals watched a 10 minute video titled *Inscapes* while undergoing fMRI. The *Inscapes* is a nonverbal, nonsocial and non-narrative computer-generated animation of shifting shapes^60^. We parcellated the brain data into the 268-ROI Shen atlas and extracted their resting-state FC matrix by calculating the Fisher-*z* transformed Pearson’s correlation between the time courses of all possible pairs of ROIs. We used the imagery CPMs, trained in the CAT dataset, to predict z-scored imagery ratings from the twins’ rsFC patterns. When the trained imagery model is applied to this independent sample, it extracts the strength of the high- (705 edges) and low-imagery network (507 edges; **Fig. 4**), applies the learned non-linear transformation, and predicts to what extent each individual thinks in the form of images on the standardized (z-scored) scale. We adopted a conservative permutation testing procedure, where we compared the actual differences in predicted imagery between the two individuals to a null distribution generated by randomly redistributing the selected edges in the imagery network across the brain while retaining the brain functional architecture before applying the model (10,000 iterations; **Suppl. Fig. 8**).

### Connectome-based classification of thought topics

We built connectome-based classification models to classify the different topics people think about at rest. Trials with more than 3 missing nodes and/or fewer than 12 available TRs after motion censoring were excluded. We further excluded trials where the speech data was not decodable or missing, or if the speech was agreed upon to belong to “Other topics” by the annotators. The 38 participants who had more than 20 valid trials were included in the analysis.

We took a leave-one-subject-out cross-validation approach. In each round of cross-validation, one participant was held-out for model testing whereas all valid trials from the remaining 59 participants were included for model training. As the topics have imbalanced numbers, in the training set, we oversampled infrequent topics using SMOTE (imblearn.over_sampling.SMOTE, https://imbalanced-learn.org/stable/references/generated/imblearn.over_sampling.SMOTE.html) and fit a non-linear support vector classification model (sklearn.svm.SVC, https://scikit-learn.org/stable/modules/generated/sklearn.svm.SVC.html, kernel = “rbf”) to classify the topics from brain connectomes. The classification accuracy of each cross-validation round was assessed as the proportion of correctly predicted trials among all valid trials in the held-out participant. Model performance was calculated as the averaged classification accuracy across all cross-validation rounds. For statistical significance testing, we compared the model performance to a null distribution generated by testing the model predicted topics on the randomly shuffled annotator-rated topics (10,000 iterations). We assumed a one-tailed significance test, with *p* = (1 + number of null *r* values ≥ empirical *r*) / (1 + number of iterations).

We additionally used balanced accuracy to correct for the imbalance between thought topics. In each round of cross-validation, we computed the balanced accuracy as the averaged recall across the eight thought topics:

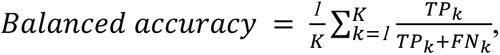

where K = number of thought topics (8), TP = true positives, and FN = false negatives. The distribution of balanced accuracy was compared to the chance level (1/8) using bootstrapping (10000 iterations).

### Canonical correlation analysis (CCA) linking model-predicted thoughts with behavior in the Human Connectome Project (HCP)

#### HCP data

We used the 1200 Subjects Release (S1200), which includes the behavioral and 3T MRI imaging data from 1206 healthy young adults. We excluded subjects without structural images or resting-state scans and further removed resting-state runs with mean frame-to-frame head displacement (FD) > 0.2 mm^121^. This gave us the final sample of 908 subjects with at least one usable resting-state run. We calculated FC patterns from the resting-state fMRI data and averaged the FC matrices within subjects if they had multiple runs.

We applied the dimensional models to the averaged rsFC pattern of each subject and generated predictions of the 9 thought dimensions. In addition, we trained a connectome-based topic classification model to classify 7 topics (movies and other topics excluded). Movies were excluded because the HCP participants did not watch the same movies as in our dataset (i.e., the CAT dataset), and other topics were excluded for clean training data. We applied the topic model to the rsFC patterns and included the predicted proportions of each topic as part of the thought matrix to reflect thought content. This resulted in a 908 × 16 matrix of model-predicted thought patterns at rest (T_1_). For measures of individual differences, we followed the exact exclusion criteria and thus kept the same set of 158 variables (from the original 478 variables) as Smith and colleagues. This gave us a 908×158 matrix of behavioral measures (S_1_). Please refer to **Suppl. Table 9** for a full list of 158 behavioral variables.

We followed the analytical procedures described in Smith et al^15^ and did the following steps to process the two matrices before feeding into the CCA model: (a) We applied a rank-based inverse Gaussian transformation to each of the columns in S_1_ and T_1_ to reduce the impact of outliers. This resulted in S_2_ and T_2_. (b) We deconfounded the S_2_ and T_2_ by regressing out the demeaned and squared measures of 17 confound variables (e.g., acquisition reconstruction software version), resulting in S_3_ and T_3_. The 17 confound variables were also identical to the ones used by the original authors. (c) We performed a principal component analysis (PCA) to reduce dimensionality for both matrices to avoid model overfitting, where we accounted for missing data by first computing the covariance matrix for subjects on a pairwise basis. For each pair of subjects, we excluded any missing data for either subject. We then projected this approximate covariance matrix onto the closest positive-definite matrix using nearestSPD (https://www.mathworks.com/matlabcentral/fileexchange/42885-nearestspd). We performed an eigenvalue decomposition (principal component analysis) to estimate the top 20 subject-wise behavioral measure eigenvector (S_4_) and the top 9 subject-wise thought eigenvector (T_4_). The choice of 9 and 20 PCs was arbitrary, however, to ensure robustness of the results, we replicated this effect irrespective of the combination of numbers of thought (ranging from 2 - 16) and behavior (10, 15, 20, 25, 30 and 35) PCs (**Suppl. Fig. 16**).

The goal of CCA is to assess the link between the transformed behavior and predicted thought matrices (S_4_ and T_4_) by finding the transformation matrices (A and B) to transform T_4_ and S_4_ to matrices of canonical weights (U and V) of the maximum correlation (https://www.mathworks.com/help/stats/canoncorr.html). We estimated 9 CCA modes, among which only the first mode significantly related predicted thoughts with behavior (*r* = .317, corrected-*p* < .001). In this first mode, A_1_ and B_1_ represent the transformation matrices and U_1_ and V_1_ represent the derived subject weights respectively from S_4_ and T_4_. The correlation between U_1_ and V_1_ is 0.309. We compared the *r*-value to a null distribution to assess significance. As there are familially-related subjects in the HCP dataset, we retained the family structure while shuffling the behavior matrix before fitting a CCA model (iteration = 10,000). We assumed a one-tail significance test and corrected the *p*-values for multiple comparisons across the 9 estimated modes.

To understand which variables in the original thoughts and behavior matrices are most strongly associated with this only identified CCA mode, we calculated the weights vectors by correlating U_1_ and V_1_ against the original behavioral measures (S_1_) and predicted thought (N_1_) matrices. This method of mapping the CCA mode to the original data matrices enabled us to estimate the same test statistics for excluded behavioral measures as well. See **Suppl. Table 10** for the sets of behavioral variables with the strongest associations (*r* > .25). See https://humanconnectome.org/storage/app/media/documentation/s500/HCP_S500_Release_Reference_Manual.pdf for the data organization and descriptions of the behavioral variables. See https://wiki.humanconnectome.org/docs/assets/HCP_S1200_DataDictionary_Oct_30_2023.csv for detailed descriptions of how these behavioral variables were defined and measured.

The identified CCA mode exhibited a positive-negative pattern of population covariation, such that favorable traits are on the positive side of the spectrum and adverse traits are on the negative side (**Suppl. Table 10**). In addition to the authors’ subjective evaluation on the valence of the variables, we utilized a pre-trained large language model–GPT4o–to rate the favorability. For each variable, GPT4o reads its definition from the above data dictionary file, interprets its semantic meanings, and rates on a 1-9 scale how favorable a higher value on this measure is. The exact prompt to perform this rating task is “Please read the attached .csv file. For each variable, please learn its semantic meanings from the definition under the ‘descriptions’ column. Explain very concisely what a higher value on the measure of this variable means, and rate on a 1-9 scale how favorable a higher value on the measure of this variable is.” Each variable was assessed independently to avoid mutual contextual influences. Some variables are not evaluable on this favorability scale, including supplied measures from the T1-weighted structural brain analysis with FreeSurfer (e.g., columns, thickness and surface areas of brain regions) and certain demographic measures that GPT4o could not provide favorability ratings on (e.g., subject ID, sex, race). For these variables, we assigned ratings of 5 to indicate that they are neither favorable nor adverse and plotted them in light gray in **Suppl. Fig. 14b**.

To investigate whether ongoing thoughts at rest contribute to the significant CCA fit above and beyond rsFC alone, we conducted a more conservative permutation testing. We estimated null distributions of CCA fit by applying null thought models, trained to predict thought dimensions and topics that were shuffled across trials within subjects from their rsFC patterns. We assumed an one-tail significance test to examine whether the CCA fits with real thought models were significantly better than the null distribution. To ensure robustness of these results, we again repeated this permutation on combinations of varied numbers of thought and behavior PCs. The CCA fit using models trained on actual (non-permuted) thought reports remains robust and consistently performed better than null (permuted) thought models across parameter settings (**Suppl. Fig.16**), suggesting that spontaneous thoughts during rest contain meaningful information about individual differences in behavior and traits, beyond what is captured by rsFC alone.

## Data availability

The raw fMRI data of our Chicago Attention and Thoughts (CAT) dataset is now private on OpenNeuro https://openneuro.org/datasets/ds006515 and will be publicly available upon publication. We created an anonymous review link to share this dataset with reviewers for anonymous access. The behavioral responses in the annotated rest task will be available on Github: https://github.com/jinke828/rest_thoughts/data/. The transcriptions of each individual’s spontaneous speech will not be open sourced due to privacy concerns. We used the 1200 Subjects release from the Human Connectome Project, which is available at https://www.humanconnectome.org/study/hcp-young-adult. Our analysis includes restricted data, the access to which can be applied here: https://www.humanconnectome.org/study/hcp-young-adult/document/restricted-data-usage.

## Code availability

The fMRI data preprocessing, main analysis and figure plotting code, the trained model weights as well as a step-by-step instruction to run these code to full replicate the key findings in this paper will be publicly available on Github: https://github.com/jinke828/rest_thoughts/code/ when the paper gets published. We utilized the canonical correlation analysis code provided by Smith and colleagues, as referenced here: https://www.fmrib.ox.ac.uk/datasets/HCP-CCA/.

## Acknowledgments

We thank the MRI Research Center at the University of Chicago (RRID: SCR_024723) for assisting with data collection. We thank Jadyn Park for sharing code to preprocess eye-tracking data, and Samantha Bertschi for annotating the thought topics. We thank the following groups and individuals for their helpful comments and suggestions on the project: the Bainbridge, Leong, Rosenberg, Bakkour (BLRB) community at the University of Chicago, the joint CogNeuro lab meeting (PIs: Marvin Chun, Nick Turk-Browne, Kia Nobre, Gregory McCarthy, Samuel McDougle and Elizabeth Goldfarb) at Yale University, the Functional Imaging & Naturalistic Neuroscience Lab (PI: Emily Finn) at Dartmouth College, Mina Kwon, Jeongjun Park and Jihyun Hur. Our work was supported by the University of Chicago Social Sciences Division, the University of Chicago Neuroscience Institute, and high-performance computing resources provided by the University of Chicago Research Computing Center and Yale Center for Research Computing.

## Author contributions

Conceptualization: J.K., T.A.C., M.D.R.

Methodology: J.K., T.A.C., A.C., H.S., M.M.C, Y.C.L., M.D.R.

Data collection: J.K., A.C., Z. Z., T. M., L. S.

Data curation, Formal analysis, & Visualization: J.K.

Supervision: M.D.R., Y.C.L., M.M.C

Funding acquisition: M.D.R.

Writing—original draft: J.K., M.D.R.

Writing—review & editing: All authors

All the authors approved the final manuscript for submission.

