## Supplementary information for "Ongoing thoughts at rest reflect functional brain organization and behavior"

Jin Ke *et al.*

**This PDF file includes:**

Figs. S1 to S16

Tables. S1 to S12

### Supplementary Figures

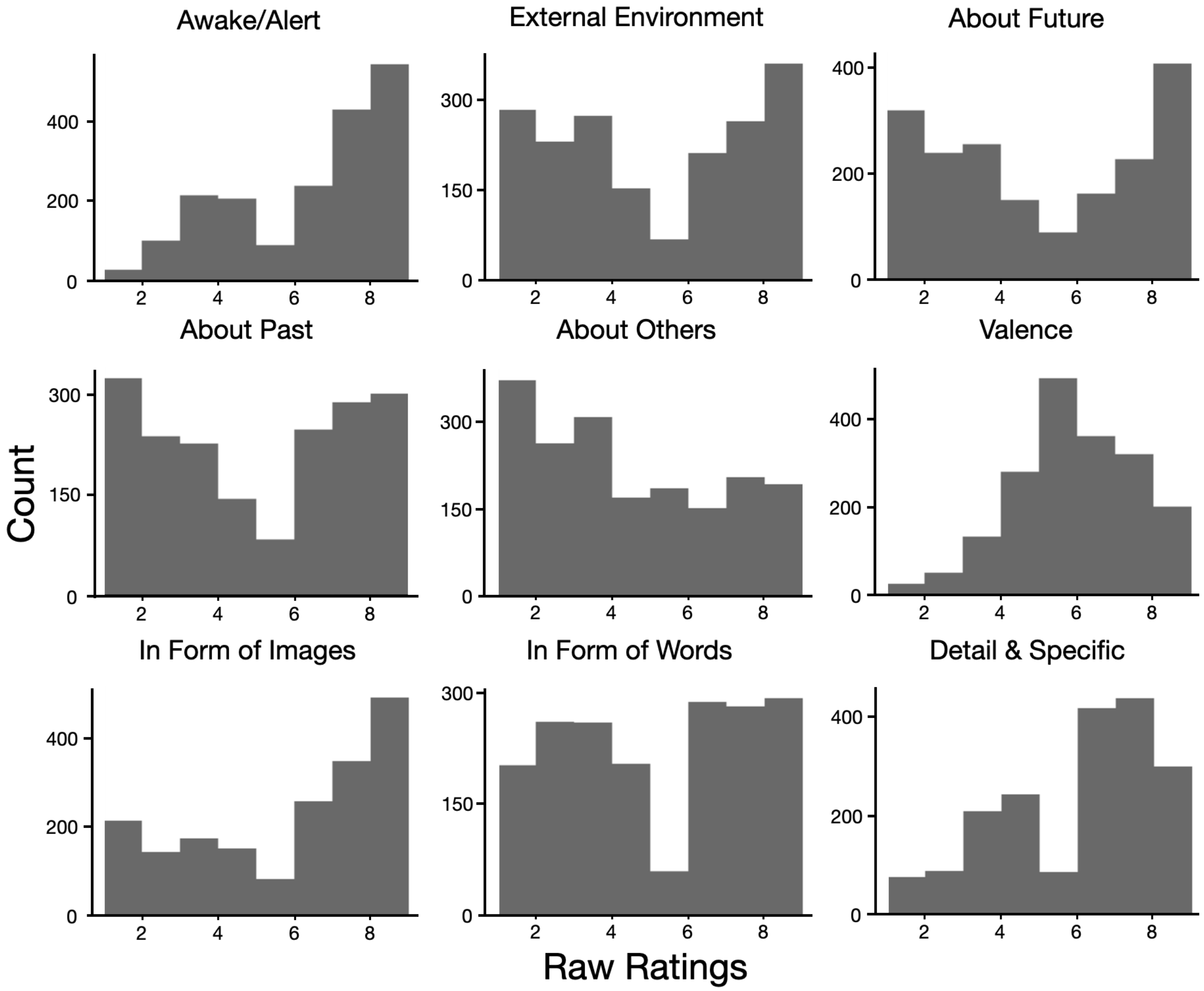


**Supplementary Figure 1.** Histograms of the raw subjective ratings of ongoing thoughts during rest for the 9 thought dimensions.

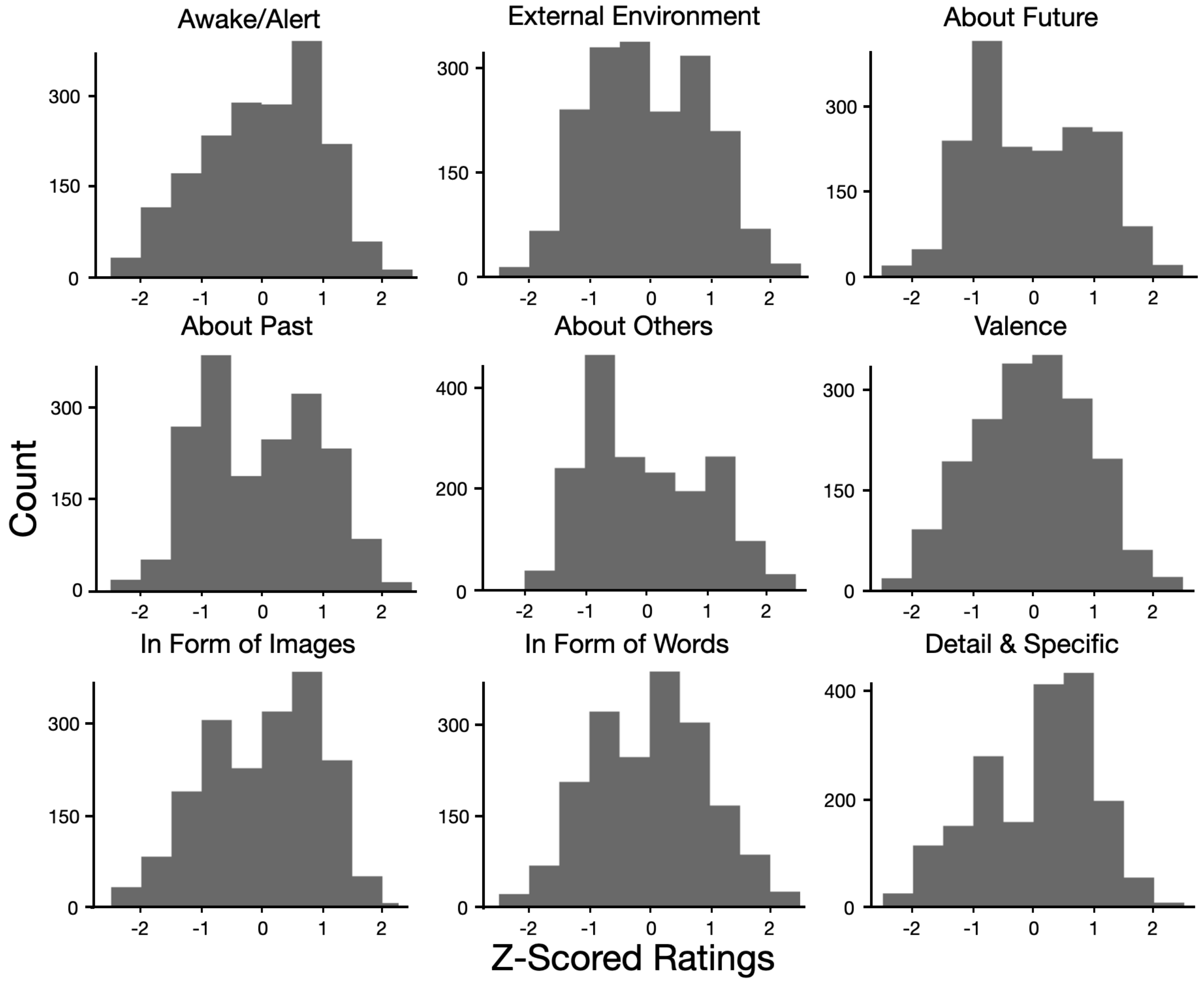


**Supplementary Figure 2.** Histograms of subjective ratings of ongoing thoughts during rest for the 9 thought dimensions. The ratings were z-normalized within individuals (to control for between-participant differences in using the slider bar) and within dimension (to control for within-participant disparity in using the slider bar differently for different dimensions).

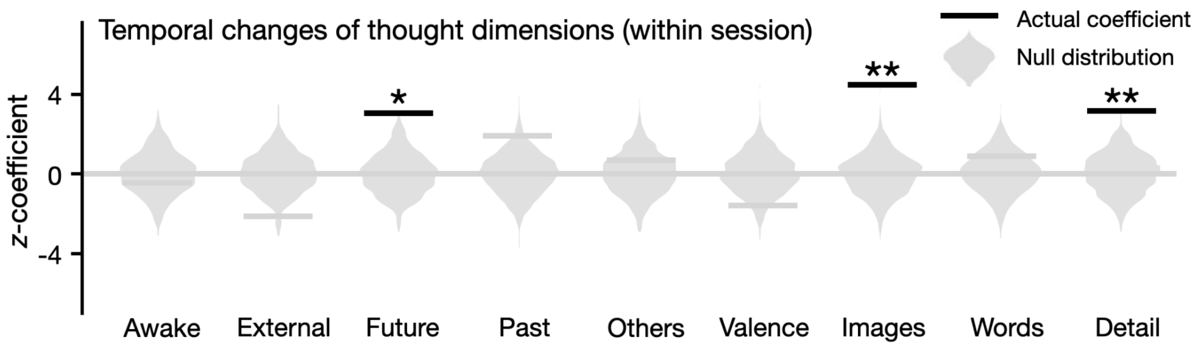


**Supplementary Figure 3.** *Z*-coefficients from nested mixed-effects models predicting ratings on each dimension by trial number (1-16). This analysis examines how dimensions of ongoing thought change across trials within scan sessions. Positive coefficients indicate increasing ratings, negative indicate decreasing ratings. Violin plots represent null distributions from shuffled ratings (**Bonferroni-corrected $p < .01$). Across the 16 trials, participants think more about the future ($\beta = .016$, *s.e.* = .005, $z = 3.042$, corrected-$p = .018$), more in the form of images ($\beta = .023$, *s.e.* = .005, $z = 4.468$, corrected-$p < .001$), and more in detail ($\beta = .016$, *s.e.* = .005, $z = 3.186$, corrected-$p < .01$).

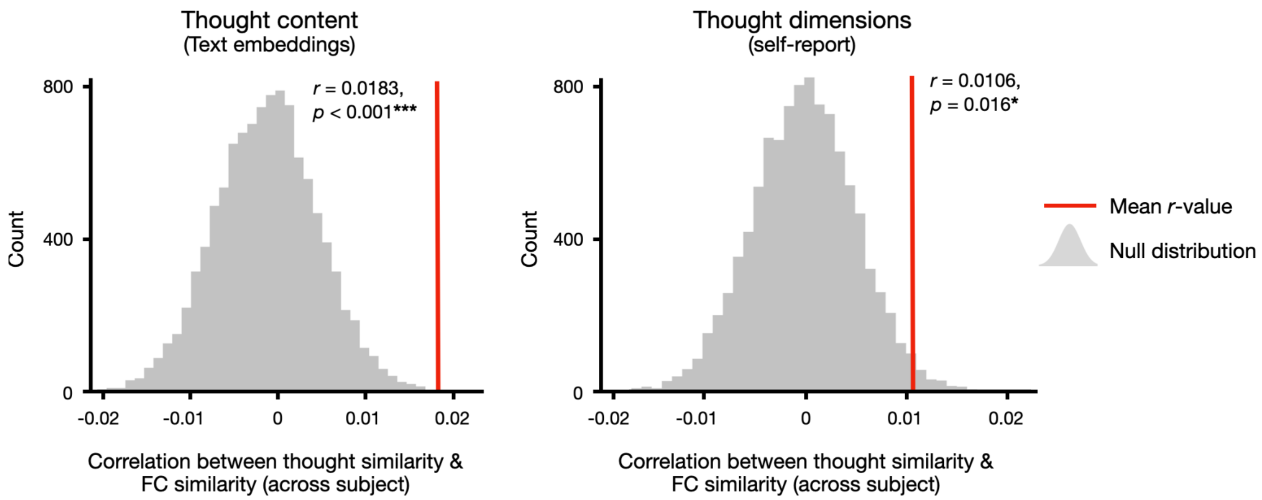


**Review Figure 4.** Between-subject trial-by-trial similarity in FC was associated with similarity in thought content (left) and dimensions (right). ***: corrected-$p < 0.001$; *: corrected-$p < 0.05$

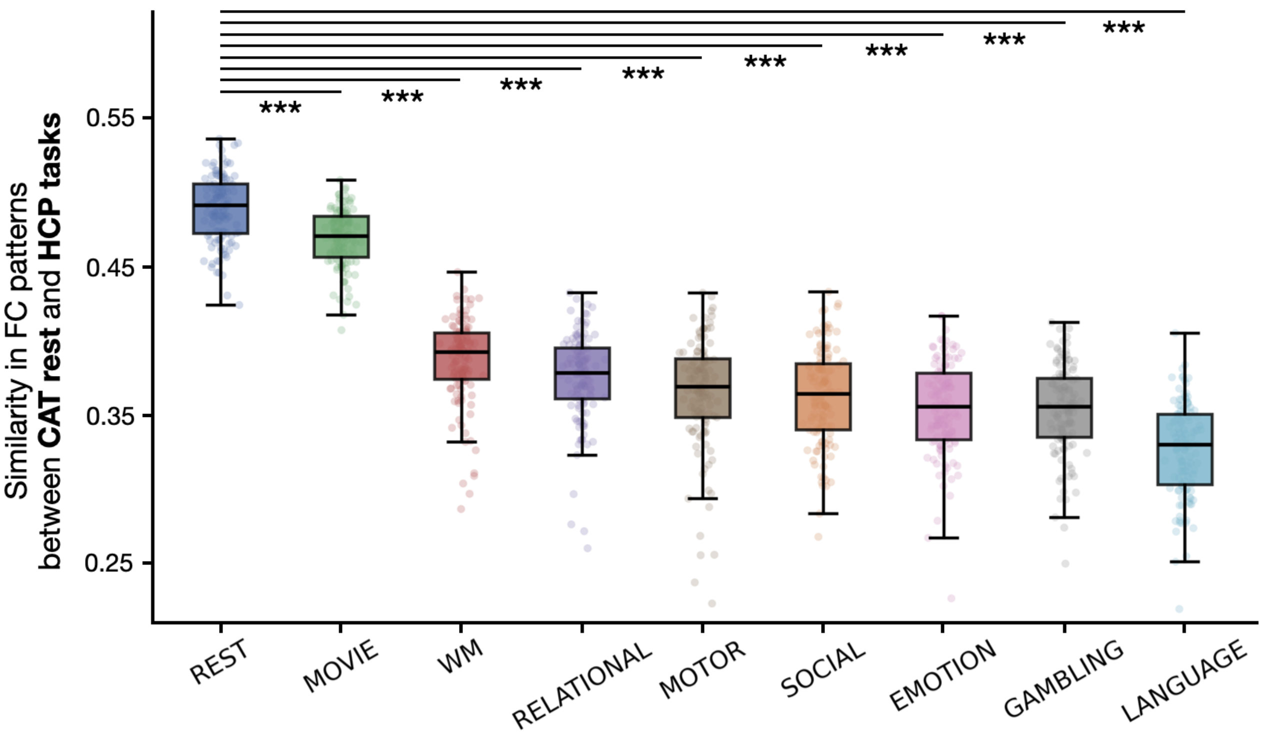


**Supplementary Figure 5.** Comparing FC patterns between CAT annotated rest and the HCP rest and tasks. For each HCP subject, correlations with every CAT subject were computed and averaged. Each dot represents the Fisher-*z* transformed mean Pearson correlation. HCP tasks are arranged by descending mean correlation. ***: corrected-$p < 0.001$

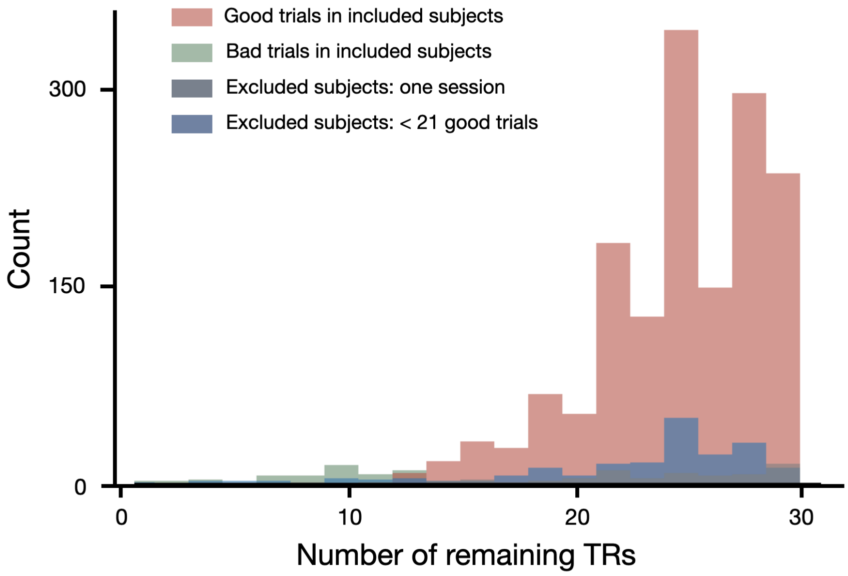


**Supplementary Figure 6.** Mean number of post–motion-censoring TRs per trial, shown separately for: excluded subjects with only a single session (gray); excluded subjects with fewer than 21 usable trials (blue); excluded trials from the 50 included subjects (green); and retained trials from the included subjects (red).

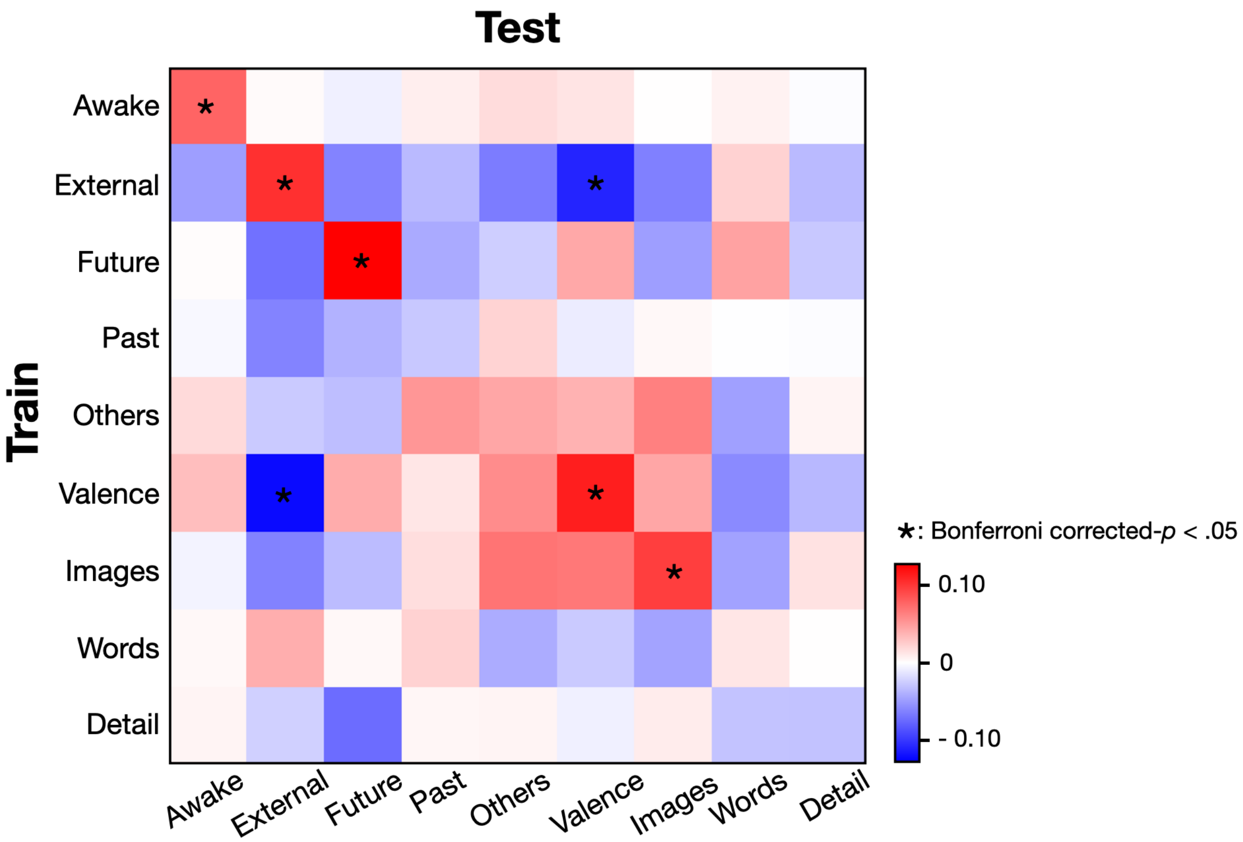


**Supplementary Figure 7.** Cross-dimensional model performance. We trained one CPM for each thought dimension and tested on all 9 thought dimensions (including itself) with leave-one-subject-out cross-validation. Each cell indicates the Pearson correlation between the model prediction (the training dimension) and participants' rating (the testing dimension). Rows represent the training dimensions and columns represent the testing dimensions. For example, the second cell on the first row means predicting ratings on external from the awake model. The diagonal represents within-dimensional prediction (**Fig. 3c**). Statistical significance was assessed with permutation, where the actual predictive accuracy was compared with a null distribution generated by shuffling the ratings within participants before testing the model. Given the ratings can be negatively correlated with each other (**Fig. 2a**), we assumed a two-tail significance test. The *p*-values were corrected for multiple comparisons across the 9 dimensions. The five significant within-dimensional CPMs exhibit higher accuracy predicting themselves than any other dimension, suggesting model specificity.

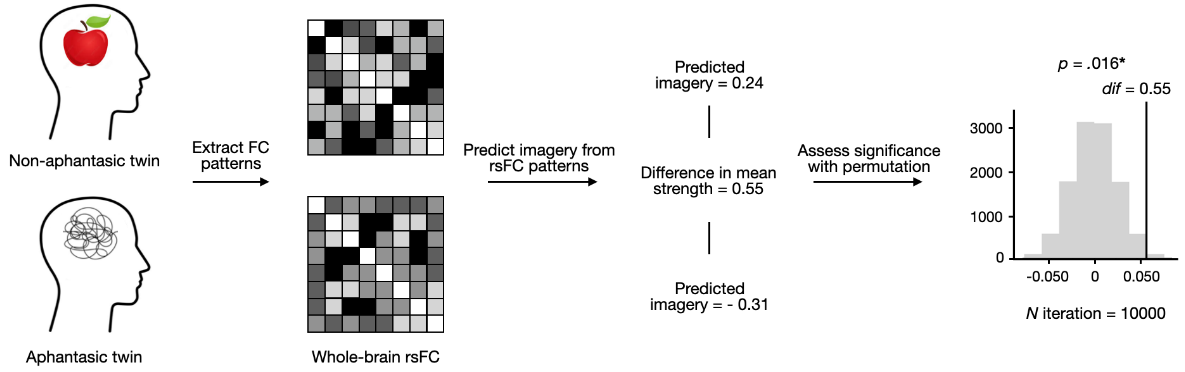


**Supplementary Figure 8.** The imagery model generalized to an external aphantasia dataset. The model predicted that an aphantasic individual showed less evidence of thinking in the form of images than their identical non-aphantastic twin. Two twins, one with aphantasia and the other without, underwent fMRI scans watching a non-narrative movie *Inscapes*. We extracted their whole-brain functional connectivity patterns and predicted to what extent they thought in the form of images with our identified imagery network. We assessed the statistical significance of the difference in the predicted imagery score between the two individuals by comparing it to a null distribution, generated by shuffling the position of selected FC features in the imagery network while retaining the brain functional architecture (10000 iterations). *: $p < .05$

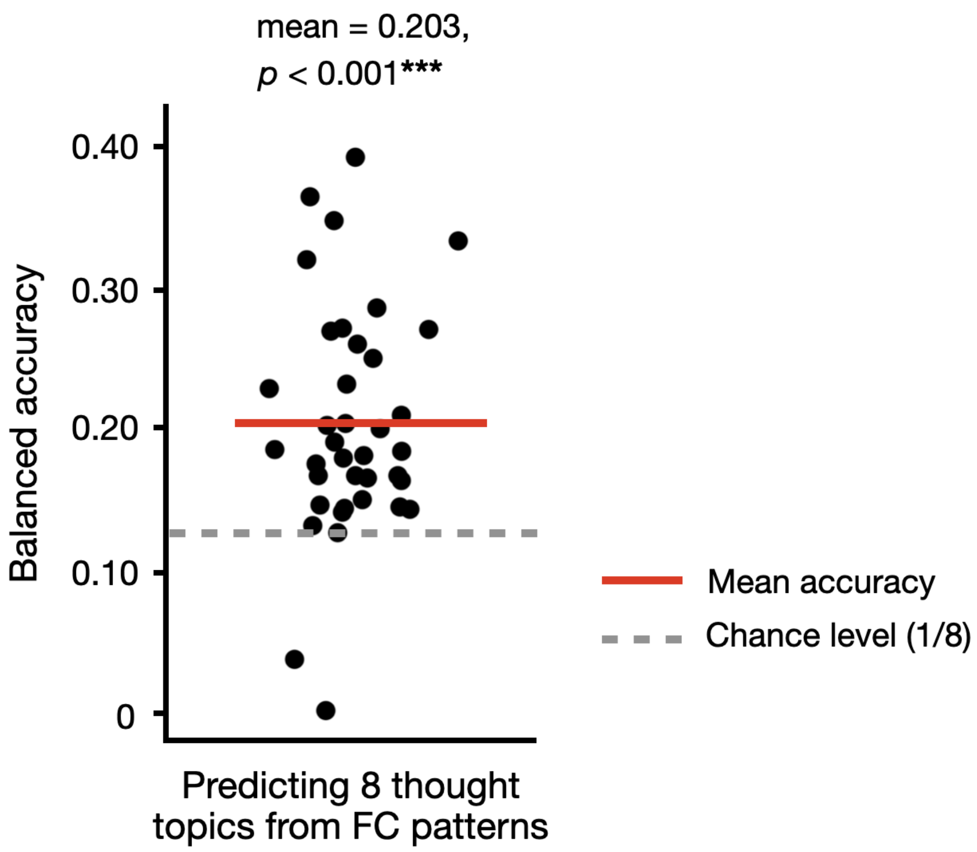


**Supplementary Figure 9.** The Balanced accuracy of the connectome-based classification models on thought topics. ***: $p < 0.001$

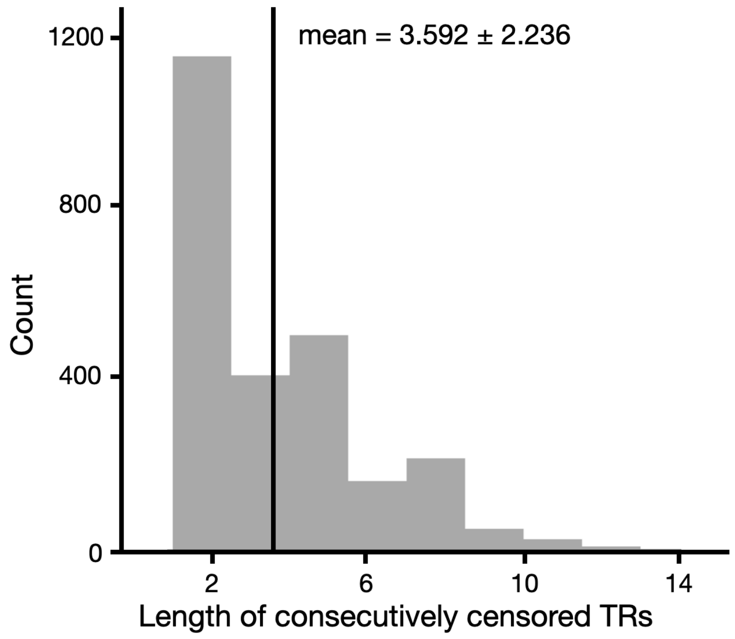


**Supplementary Figure 10.** Empirical assessment of spin-history effects in the CAT annotated-rest dataset. We plotted the distribution of consecutive censored TRs and found that, on average, approximately 2.6 additional TRs were censored following an initial censored volume.

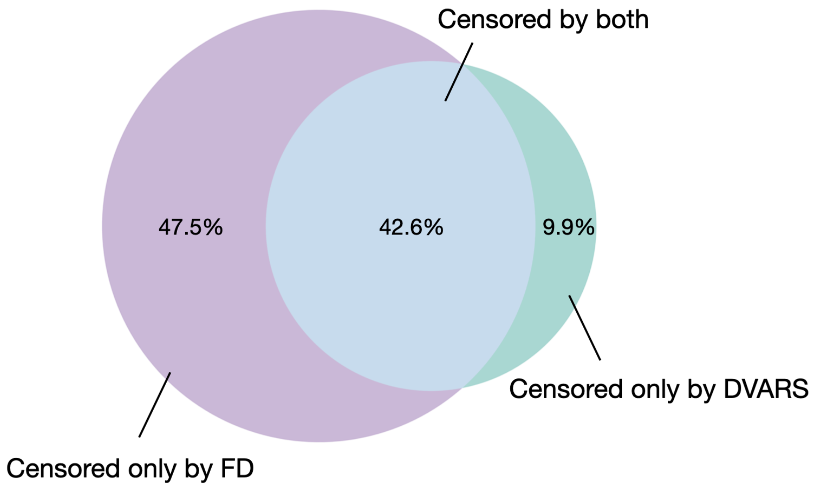


**Supplementary Figure 11.** Proportion of censored TRs across motion categories: censored solely due to frame-wise displacement > 0.25 mm, solely due to DVARS ($p < 0.05$), and censored by both criteria.

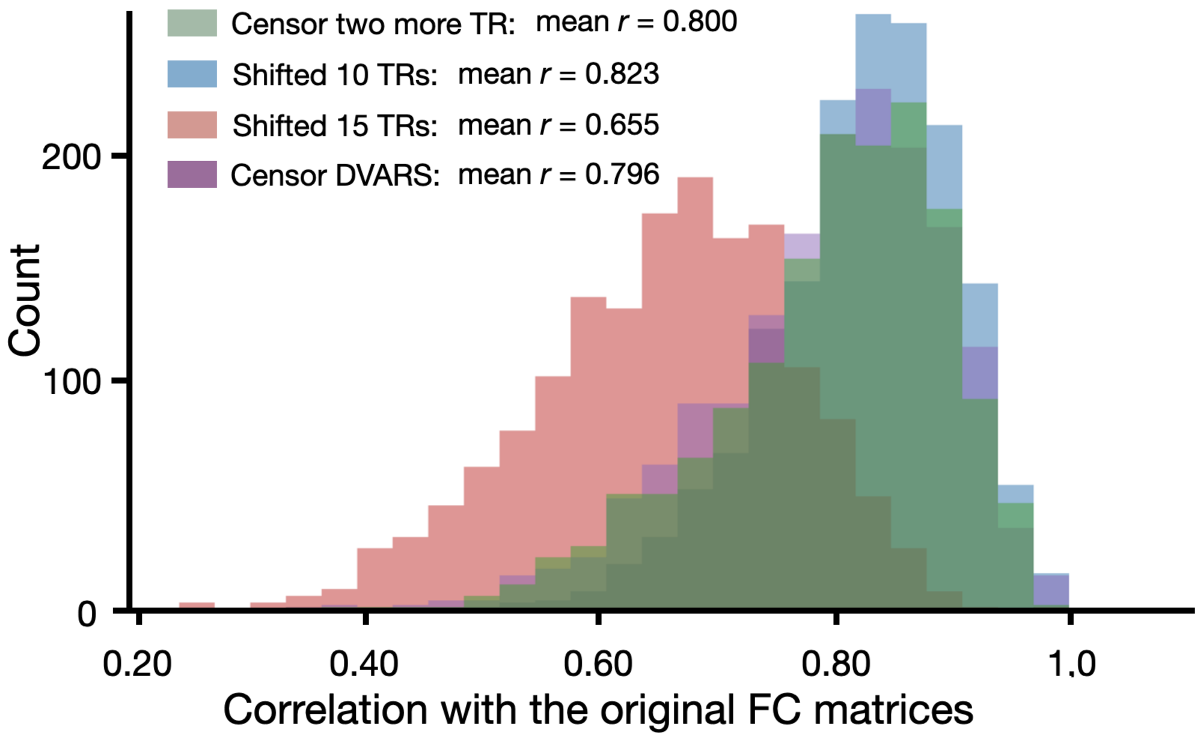


**Supplementary Figure 12.** Correlations between the original rsFC patterns and multiple control variants. To assess potential hemodynamic lag effects, we temporally shifted the fMRI time series by 10 TRs (blue) and 15 TRs (red) prior to rsFC estimation, censored two additional TRs following each motion-censored frame (green), and applying additional DVARS-based censoring (purple).

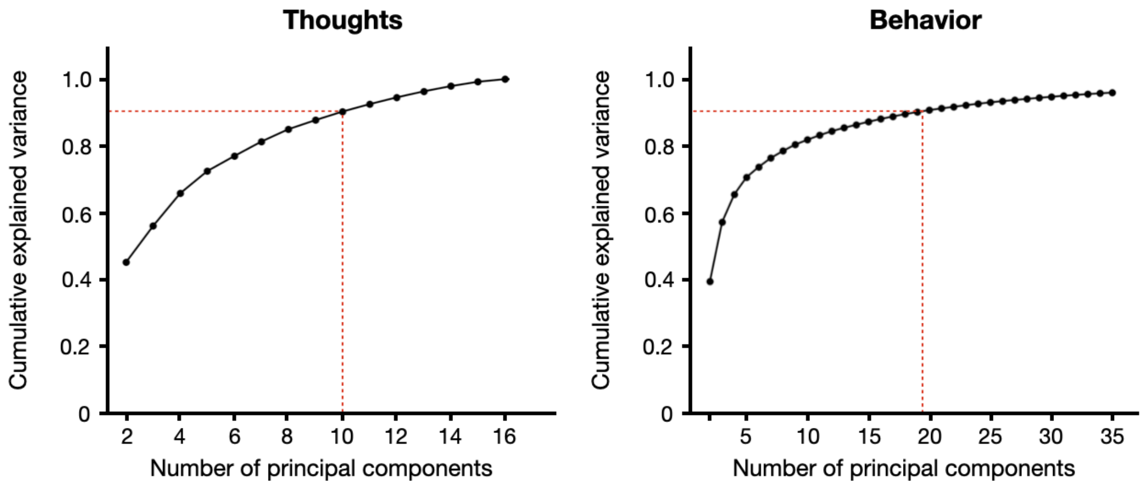


**Supplementary Figure 13.** Cumulative explained variance as a function of the number of principal components for thought measures (left) and behavioral measures (right). Thought dimensions required 9 principal components to reach 90% explained variance, whereas behavioral measures required 20.

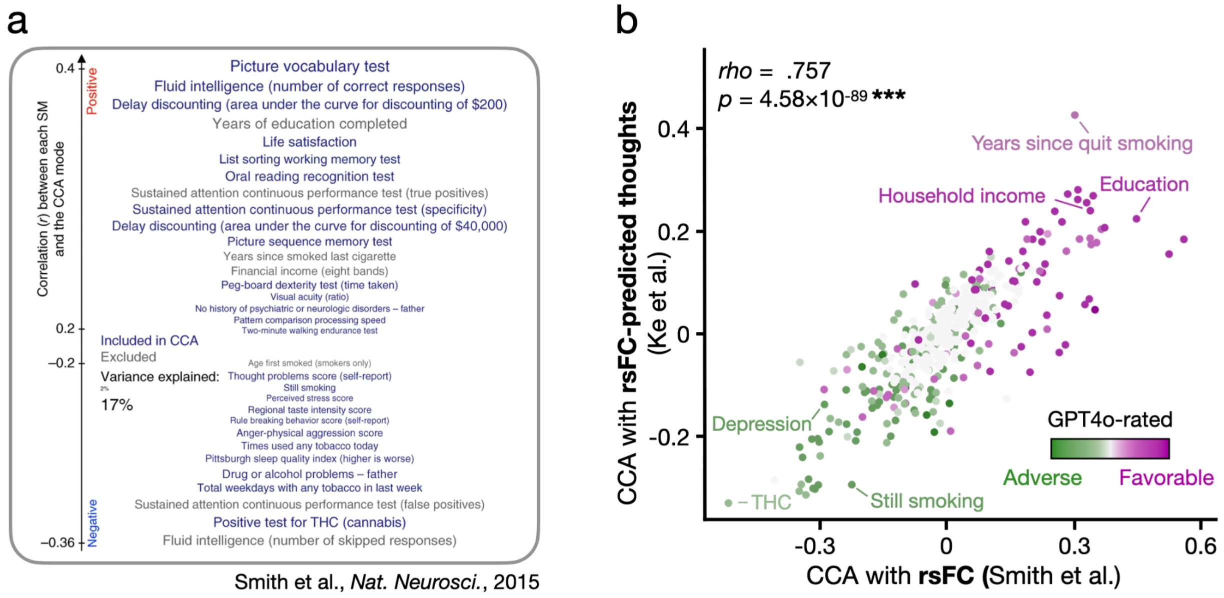


**Supplementary Figure 14.** (**a**) Top behavioral associations from Smith et al., who linked the same measures with rsFC with CCA. **(b)** Comparison on variable weights between this study and Smith et al. The y-axis represents variable weights in the current study, linking individual differences measures with model-predicted thoughts; the x-axis represents variable weights of Smith et al, linking the same individual differences measures with rsFC. Each dot represents a behavioral variable, colored by GPT4o-rated favorability based on its interpretations on the variable definition (See *Methods*).

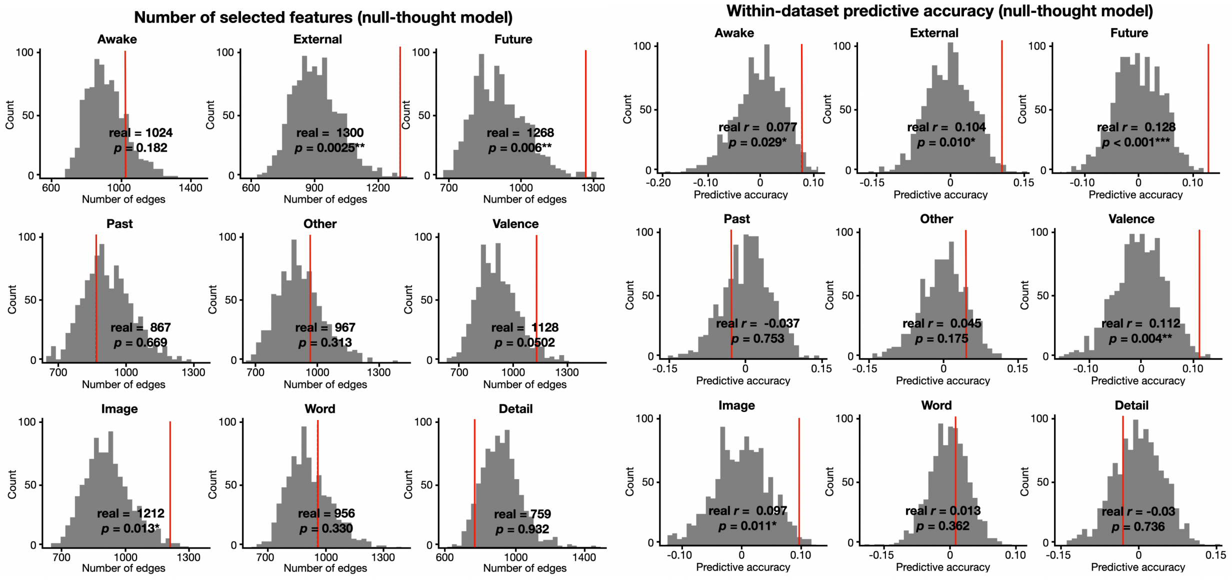


**Review Figure 15**. Distributions of the number of selected features (left) and within-dataset predictive accuracy (right) of the null-thought models. These null models were built by shuffling the thoughts dimensions and topics across trials within subjects.

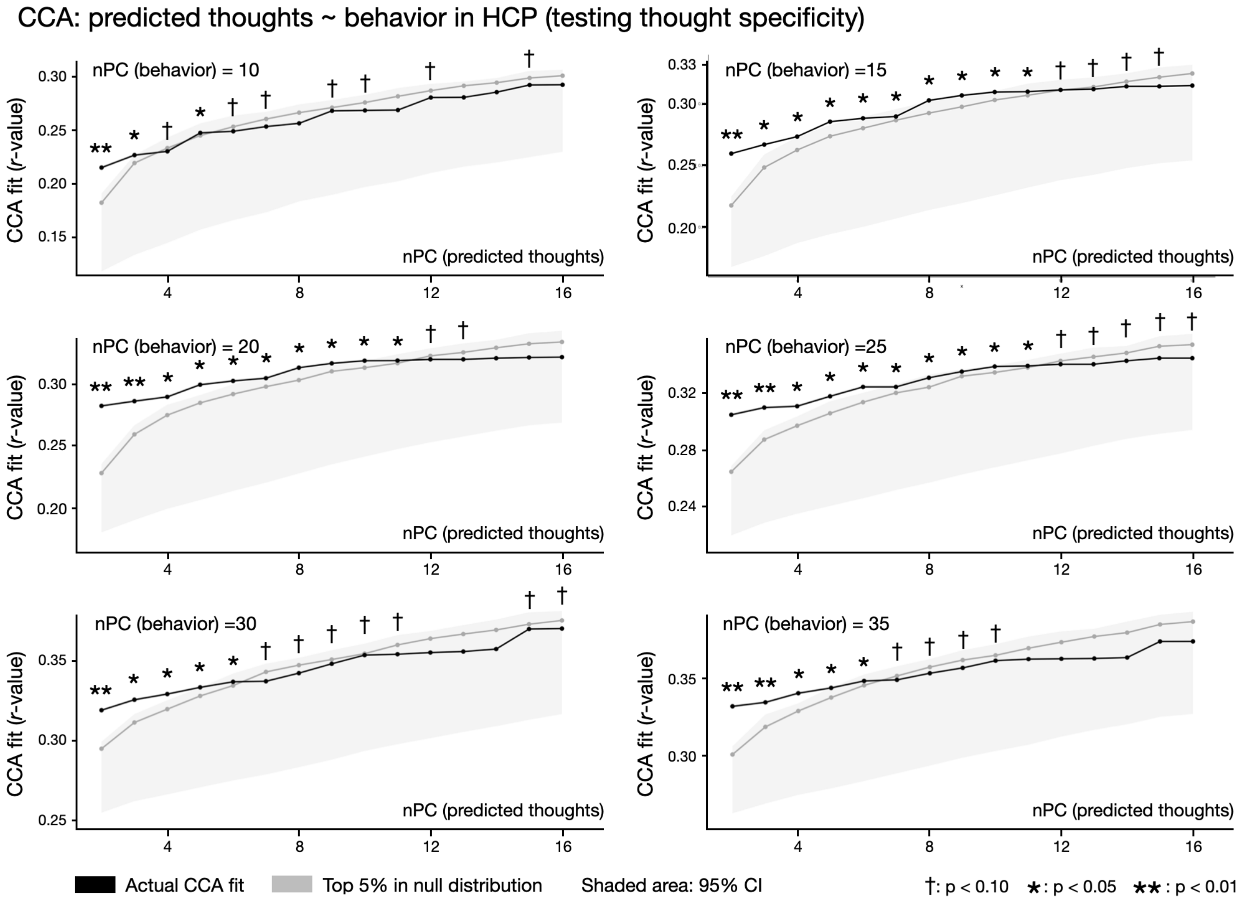


**Supplementary figure 16.** Predicted ongoing thoughts at rest show unique contributions to the CCA fit above and beyond rsFC alone. The black line shows the *r*-value of the primary CCA mode, which is identical to the black line in **Fig. S7**. The gray line shows the 95th percentile of the null distribution. The null thought models were built in our CAT dataset to predict behavioral ratings/topics shuffled within participants from rsFC patterns (iterations = 1000). The shaded area indicates 95% confidence interval of the null distribution. We note that the upper boundary thus denotes the 97.5th percentile of the null distribution. Statistical significance was assessed with a one-tail test against the above-described null distribution.

### Supplementary Tables

**Table S1. The mean and standard deviation of ratings within- and between-participants.**

| Dimensions | Mean | Standard deviation (within-participant) | Standard deviation (across-participant) |
|---|---|---|---|
| Awake | 6.064 | 1.453 | 1.491 |
| External | 4.746 | 2.267 | 1.363 |
| Future | 4.665 | 2.403 | 1.250 |
| Past | 4.581 | 2.328 | 1.164 |
| Other | 3.995 | 2.277 | 0.782 |
| Valence | 5.431 | 1.443 | 0.679 |
| Image | 5.447 | 2.077 | 1.441 |
| Word | 4.735 | 1.925 | 1.512 |
| Detail | 5.542 | 1.800 | 1.021 |

Note: To calculate within-participant standard deviation, we calculated the standard deviations of ratings across trials for each participant and averaged the values across participants. To calculate across-participants standard deviation, we took the mean of the ratings for each participant and calculated the standard deviation of this mean rating distribution.

**Table S2. The definitions of each thought topic. Five annotators rated the transcriptions of free speech into topics based on these definitions as the instructions.**

| Topic | Descriptions |
|---|---|
| Current environment | Thoughts about the current environment (i.e., inside the scanner), including thoughts about the annotated rest task itself |
| Body sensations and introspections | Thoughts or feelings about their own body or bodily sensations as well as awareness of their own thoughts and other internal mental processes |
| Movies | Thoughts about the naturalistic stimuli they watched and/or listen to in the scanner (i.e., pie podcast, croissant video, *North by Northwest* clip, and cake video) |
| Social | Thoughts primarily about other people |
| Future planning | Thoughts about making short-term or long-term plans and/or things that will happen in the future when such thoughts do not clearly fit in the social category |
| Past memory | Thoughts related to what people have experienced in the past when such thoughts do not clearly fit in the social category |
| Obligations | Thoughts related to study, work or things they have to do that don't clearly fall into the future planning or past memory categories |
| Zoning out | When people reported they were zoning out or not thinking about anything in particular |
| Other topics | Thoughts that do not categorize into any of the other 8 topics |

**Table S3. Temporal changes of thought dimensions when controlling for between-trial autocorrelations.**

| Dimensions | Original modeling (Fig. 2b) | Regress autocorrelation (9 dimensions) | Regress raw difference (1 dimension) | Regress absolute difference (1 dimension) |
|---|---|---|---|---|
| Awake | $\beta$ = -3.440, $p$ = 0.009** | $\beta$ = -2.927, $p$ = 0.027* | $\beta$ = -2.884, $p$ = 0.036* | $\beta$ = -3.617, $p$ = 0.009** |
| External | $\beta$ = -5.905, $p$ < 0.001*** | $\beta$ = -4.642, $p$ < 0.001*** | $\beta$ = -5.721, $p$ < 0.001*** | $\beta$ = -4.552, $p$ < 0.001*** |
| Future | $\beta$ = 2.342, $p$ = 0.171 | $\beta$ = 1.757, $p$ = 0.711 | $\beta$ = 2.685, $p$ = 0.063 | $\beta$ = 1.739, $p$ = 0.738 |
| Past | $\beta$ = 4.200, $p$ < 0.001*** | $\beta$ = 3.401, $p$ = 0.001*** | $\beta$ = 5.539, $p$ < 0.001*** | $\beta$ = 3.310, $p$ = 0.009** |
| Other | $\beta$ = 0.704, $p$ = 1.0 | $\beta$ = 0.611, $p$ =1.0 | $\beta$ = 0.820, $p$ = 1.0 | $\beta$ = 0.596, $p$ = 1.0 |
| Valence | $\beta$ = -0.839, $p$ = 1.0 | $\beta$ = -1.179, $p$ = 1.0 | $\beta$ = -1.127, $p$ = 1.0 | $\beta$ = -1.147, $p$ = 1.0 |
| Image | $\beta$ = 4.991, $p$ < 0.001*** | $\beta$ = 3.351, $p$ = 0.009** | $\beta$ = 6.152, $p$ < 0.001*** | $\beta$ = 3.356, $p$ = 0.009** |
| Word | $\beta$ = -1.315, $p$ = 1.0 | $\beta$ = -0.526, $p$ = 1.0 | $\beta$ = -0.500, $p$ = 1.0 | $\beta$ = -0.831, $p$ = 1.0 |
| Detail | $\beta$ = 0.260, $p$ = 1.0 | $\beta$ = 0.682, $p$ = 1.0 | $\beta$ = 1.623, $p$ = 0.945 | $\beta$ = 0.883, $p$ =1.0 |

*: $p$ < 0.05, **: $p$ < 0.01, ***: $p$ < 0.001

**Table S4.** Summary of subjects and trials excluded from the CPM analyses, along with the distribution of remaining TRs per trial after motion censoring.

| | Reason for excluding | *N* dropped/ included | Mean number of remaining TR (per trial) |
|---|---|---|---|
| Excluded subjects (*N* = 10) | Did only one scan session | N = 4 (64 trials) | 22.65 ± 5.73 |
| | < 21 good trials in total | N = 6 (92 trials) | 23.92 ± 5.21 |
| Included subjects (*N* = 50) | < 12 TRs available | 61 trials | 8.72 ± 3.19 |
| | — | 1531 trials | 24.73 ± 3.85 |

**Table S5. Performance of connectome-based predictive models of thought dimensions**

| **Dimension** | ***r*** | ***p* (uncorrected)** | ***p* (corrected)** | **MSE** | ***p* (uncorrected)** | ***p* (corrected)** | **$R^2$** | ***p* (uncorrected)** | ***p* (corrected)** |
|---|---|---|---|---|---|---|---|---|---|
| Awake | 0.077 | **0.002** | **0.021** | 1.076 | **0.002** | **0.014** | -0.071 | **0.002** | **0.014** |
| External | 0.104 | **0.0003** | **0.003** | 1.060 | **<.0001** | **0.001** | -0.043 | **<.0001** | **0.001** |
| Future | 0.128 | **<.0001** | **0.001** | 1.046 | **<.0001** | **0.001** | -0.044 | **<.0001** | **0.001** |
| Past | -0.027 | 0.839 | 1.0 | 1.153 | 0.882 | 1.0 | -0.14 | 0.880 | 1.0 |
| Other | 0.045 | **0.049** | 0.441 | 1.095 | **0.036** | 0.328 | -0.089 | **0.041** | 0.370 |
| Valence | 0.112 | **0.0002** | **0.002** | 1.038 | **0.0002** | **0.002** | -0.036 | **<.0001** | **0.001** |
| Image | 0.097 | **0.0003** | **0.003** | 1.065 | **0.0006** | **0.005** | -0.065 | **0.0005** | **0.004** |
| Word | 0.013 | 0.311 | 1.0 | 1.087 | 0.287 | 1.0 | -0.078 | 0.273 | 1.0 |
| Detail | -0.031 | 0.870 | 1.0 | 1.148 | 0.920 | 1.0 | -0.132 | 0.942 | 1.0 |

Note: Significance on each of the three evaluative matrices was assessed through permutation, where ratings of the held-out participant were shuffled before correlating with the real model predictions. Here, we presented *p*-values both before and after correction for multiple comparisons (Bonferroni) and *p*-values < 0.05 were marked in bold. We used correlation (*r*) as our main evaluation matrix as we were interested in whether our model captures the within-individual thought patterns, rather than the actual value of these ratings in each trial. Nevertheless, we reported all three matrices (*r*, MSE and *r*-squared). We note that the $R^2$ measures were negative despite being significantly above the null distribution, supporting our choice of Pearson's *r* as the main indicator of the model performance (Song et al., 2021).

**Table S6.** Replicating the key findings of this work with newly estimated rsFC patterns from variants of complementary control analysis to assess the impact of motion.

| Analysis | | Original results | Censor 2 more TR | Shift 10 TRs | Shift 15 TRs | DVARS |
|---|---|---|---|---|---|---|
| **RSA** (Fig. 3b) | Content (Between subject) | $r = 0.096$, $p < 0.001$*** | $r = 0.052$, $p = 0.016$* | $r = 0.067$, $p = 0.002$** | $r = 0.070$, $p = 0.003$** | $r = 0.075$, $p < 0.001$*** |
| | Dimensions (Between subject) | $r = 0.026$, $p = 0.141$ | $r = 0.024$, $p = 0.165$ | $r = 0.018$, $p = 0.221$ | $r = 0.005$, $p = 0.412$ | $r = 0.019$, $p = 0.199$ |
| | Content (within-subject) | $r = 0.069$, $p < 0.001$*** | $r = 0.061$, $p < 0.001$*** | $r = 0.060$, $p < 0.001$*** | $r = 0.069$, $p < 0.001$*** | $r = 0.059$, $p < 0.001$*** |
| | Dimensions (within-subject) | $r = 0.071$, $p < 0.001$*** | $r = 0.064$, $p < 0.001$*** | $r = 0.074$, $p < 0.001$*** | $r = 0.081$, $p < 0.001$*** | $r = 0.069$, $p < 0.001$*** |
| **CPMs - dimensions** (Fig. 3c) (all Bonferroni-corrected *p*) | Awake | $r = 0.077$, $p = 0.021$* | $r = 0.074$, $p = 0.045$* | $r = 0.091$, $p < 0.001$*** | $r = 0.061$, $p = 0.099$ | $r = 0.079$, $p < 0.001$*** |
| | External | $r = 0.104$, $p = 0.003$** | $r = 0.127$, $p < 0.001$*** | $r = 0.058$, $p = 0.126$ | $r = 0.094$, $p < 0.001$*** | $r = 0.060$, $p = 0.143$ |
| | Future | $r = 0.128$, $p = 0.001$** | $r = 0.164$, $p < 0.001$*** | $r = 0.131$, $p < 0.001$*** | $r = 0.109$, $p < 0.001$*** | $r = 0.104$, $p < 0.001$*** |
| | Past | $r = -0.027$, $p = 1.0$ | $r = 0.046$, $p = 0.496$ | $r = 0.040$, $p = 0.746$ | $r = 0.016$, $p = 1.0$ | $r = 0.002$, $p = 1.0$ |
| | Other | $r = 0.045$, $p = 0.441$ | $r = -0.045$, $p = 1$ | $r = 0.044$, $p = 0.369$ | $r = 0.083$, $p < 0.001$*** | $r = 0.034$, $p = 0.890$ |
| | Valence | $r = 0.112$, $p = 0.002$** | $r = 0.106$, $p < 0.001$*** | $r = 0.129$, $p < 0.001$*** | $r = 0.121$, $p < 0.001$*** | $r = 0.178$, $p < 0.001$*** |
| | Image | $r = 0.097$, $p = 0.003$** | $r = 0.105$, $p < 0.001$*** | $r = 0.164$, $p < 0.001$*** | $r = 0.121$, $p < 0.001$*** | $r = 0.142$, $p < 0.001$*** |
| | Word | $r = 0.013$, $p = 1.0$ | $r = 0.039$, $p = 0.710$ | $r = 0.014$, $p = 1$ | $r = -0.052$, $p = 1$ | $r = 0.063$, $p = 0.054$ |
| | Detail | $r = -0.031$, $p = 1.0$ | $r = 0.043$, $p = 0.539$ | $r = 0.053$, $p = 0.243$ | $r = -0.028$, $p = 1$ | $r = 0.072$, $p = 0.036$* |
| **CPM - topics** (Fig. 3d) | | accuracy = 20.3%, $p < 0.001$*** | accuracy = 19.4%, $p < 0.001$*** | accuracy = 19.6%, $p < 0.001$*** | accuracy = 19.5%, $p < 0.001$*** | accuracy = 19.3%, $p < 0.001$*** |
| **Predicting Aphantasia** (Suppl. Fig. 5) | | dif = 0.55, $p = 0.016$* | dif = 0.416, $p = 0.126$ | dif = 0.484, $p = 0.043$* | dif = 0.355, $p = 0.072$ | dif = 0.486, $p = 0.031$* |
| **CCA** (Fig. 5b) | | $r = 0.317$, $p < 0.001$*** | $r = 0.290$, $p = 0.0014$** | $r = 0.293$, $p < 0.001$*** | $r = 0.293$, $p = 0.001$** | $r = 0.291$, $p < 0.001$*** |

*: $p < 0.05$, **: $p < 0.01$, ***: $p < 0.001$

**Table S7.**. Within-subject, trial-wise correlations between mean frame-wise displacement (left) and number of TR censored with thought ratings on each of the dimensions.

**Correlating trial-to-trial frame-wise displacement (FD) with thought ratings**

| Dimensions | *r*-values | *p*-values | corrected-*p* |
|---|---|---|---|
| Awake | 0.002 | 0.930 | 1.0 |
| External | -0.048 | 0.054 | 0.489 |
| Future | 0.036 | 0.150 | 1.0 |
| Past | 0.021 | 0.398 | 1.0 |
| Other | 0.006 | 0.802 | 1.0 |
| Valence | 0.041 | 0.101 | 0.906 |
| Image | 0.024 | 0.346 | 1.0 |
| Word | -0.015 | 0.561 | 1.0 |
| Detail | -0.021 | 0.399 | 1.0 |

*n.s.* in all conditions

**Correlating trial-to-trial number of TRs (after motion censoring) with thought ratings**

| Dimensions | *r*-values | *p*-values | corrected-*p* |
|---|---|---|---|
| Awake | -0.034 | 0.146 | 1.0 |
| External | 0.009 | 0.703 | 1.0 |
| Future | 0.146 | 0.534 | 1.0 |
| Past | -0.008 | 0.721 | 1.0 |
| Other | 0.003 | 0.887 | 1.0 |
| Valence | -0.037 | 0.111 | 0.996 |
| Image | 0.021 | 0.365 | 1.0 |
| Word | 0.033 | 0.162 | 1.0 |
| Detail | 0.021 | 0.378 | 1.0 |

*n.s.* in all conditions

**Table S8.** Across-subject correlations between mean frame-wise displacement, number of TR censored, and temporal signal-to-noise ratio with CPM performance on each of the dimensions.

| Dimensions | Mean frame-wise displacement (FD) | Number of TR censored | Temporal signal-to-noise ratio (tSNR) |
|---|---|---|---|
| Awake | $r = 0.010$, $p = 0.943$, corrected-$p = 1.0$ | $r = -0.091$, $p = 0.529$, corrected-$p = 1.0$ | $r = -0.186$, $p = 0.197$, corrected-$p = 1.0$ |
| External | $r = 0.239$, $p = 0.094$, corrected-$p = 0.844$ | $r = 0.229$, $p = 0.110$, corrected-$p = 0.988$ | $r = 0.244$, $p = 0.087$, corrected-$p = 0.785$ |
| Future | $r = -0.068$, $p = 0.643$, corrected-$p = 1.0$ | $r = -0.114$, $p = 0.434$, corrected-$p = 1.0$ | $r = -0.0356$, $p = 0.808$, corrected-$p = 1.0$ |
| Past | $r = -0.117$, $p = 0.423$, corrected-$p = 1.0$ | $r = -0.114$, $p = 0.434$, corrected-$p = 1.0$ | $r = 0.104$, $p = 0.478$, corrected-$p = 1.0$ |
| Other | $r = -0.017$, $p = 0.904$, corrected-$p = 1.0$ | $r = -0.134$, $p = 0.354$, corrected-$p = 1.0$ | $r = 0.065$, $p = 0.654$, corrected-$p = 1.0$ |
| Valence | $r = -0.233$, $p = 0.104$, corrected-$p = 0.936$ | $r = -0.231$, $p = 0.106$, corrected-$p = 0.954$ | $r = 0.129$, $p = 0.373$, corrected-$p = 1.0$ |
| Image | $r = -0.058$, $p = 0.692$, corrected-$p = 1.0$ | $r = -0.104$, $p = 0.472$, corrected-$p = 1.0$ | $r = 0.069$, $p = 0.634$, corrected-$p = 1.0$ |
| Word | $r = -0.077$, $p = 0.598$, corrected-$p = 1.0$ | $r = -0.104$, $p = 0.471$, corrected-$p = 1.0$ | $r = 0.143$, $p = 0.322$, corrected-$p = 1.0$ |
| Detail | $r = -0.081$ $p = 0.576$, corrected-$p = 1.0$ | $r = -0.124$, $p = 0.392$, corrected-$p = 1.0$ | $r = -0.058$, $p = 0.688$, corrected-$p = 1.0$ |

*n.s.* in all conditions

**Table S9.** A full list of subjective measures used for canonical correlation analysis in the HCP and MBB datasets (in alphabetical order)

| Human Connectome Project (HCP) - 158 subjective measures |
| --- |
| AngAffect_Unadj AngAggr_Unadj AngHostil_Unadj ASR_Aggr_Pct ASR_Aggr_Raw ASR_Anxd_Pct ASR_Anxd_Raw ASR_Attn_Pct ASR_Attn_Raw ASR_Crit_Raw ASR_Extn_Raw ASR_Extn_T ASR_Intn_Raw ASR_Intn_T ASR_Intr_Pct ASR_Intr_Raw ASR_Oth_Raw ASR_Rule_Pct ASR_Rule_Raw ASR_Soma_Pct ASR_Soma_Raw ASR_TAO_Sum ASR_Thot_Pct ASR_Thot_Raw ASR_Totp_Raw ASR_Totp_T ASR_Witd_Pct ASR_Witd_Raw Avg_Weekday_Any_Tobacco_7days Avg_Weekday_Beer_Wine_Cooler_7days Avg_Weekday_Cigarettes_7days Avg_Weekday_Drinks_7days Avg_Weekday_Wine_7days Avg_Weekend_Any_Tobacco_7days Avg_Weekend_Beer_Wine_Cooler_7days Avg_Weekend_Cigarettes_7days Avg_Weekend_Drinks_7days Avg_Weekend_Hard_Liquor_7days Avg_Weekend_Wine_7days CardSort_AgeAdj CardSort_Unadj Correction DDisc_AUC_200 DDisc_AUC_40K Dexterity_AgeAdj Dexterity_Unadj DSM_Adh_Pct DSM_Adh_Raw DSM_Antis_Pct DSM_Antis_Raw DSM_Anxi_Pct DSM_Anxi_Raw DSM_Avoid_Pct DSM_Avoid_Raw DSM_Depr_Pct DSM_Depr_Raw DSM_Hype_Raw DSM_Inat_Raw DSM_Somp_Pct DSM_Somp_Raw EmotSupp_Unadj ER40_CR ER40ANG ER40FEAR ER40NOE ER40SAD EVA_Denom FamHist_Fath_Dep FamHist_Fath_DrgAlc FamHist_Fath_None FamHist_Moth_Dep FamHist_Moth_None FearAffect_Unadj FearSomat_Unadj Flanker_AgeAdj Flanker_Unadj Friendship_Unadj InstruSupp_Unadj IWRD_TOT LifeSatisf_Unadj ListSort_AgeAdj ListSort_Unadj Loneliness_Unadj Mars_Final Mars_Log_Score MeanPurp_Unadj MMSE_Score NEOFAC_A NEOFAC_C NEOFAC_E NEOFAC_N NEOFAC_O Num_Days_Drank_7days Num_Days_Used_Any_Tobacco_7days Odor_AgeAdj Odor_Unadj PainInterf_Tscore PercHostil_Unadj PercReject_Unadj PercStress_Unadj PicSeq_AgeAdj PicSeq_Unadj PicVocab_AgeAdj PicVocab_Unadj PMAT24_A_CR PosAffect_Unadj ProcSpeed_AgeAdj ProcSpeed_Unadj PSQI_Score ReadEng_AgeAdj ReadEng_Unadj Sadness_Unadj SCPT_SEN SCPT_SPEC SelfEff_Unadj SSAGA_Agoraphobia SSAGA_Alc_12_Drinks_Per_Day SSAGA_Alc_12_Frq SSAGA_Alc_12_Frq_5plus SSAGA_Alc_12_Frq_Drk SSAGA_Alc_12_Max_Drinks SSAGA_Alc_Age_1st_Use SSAGA_Alc_D4_Ab_Dx SSAGA_Alc_D4_Ab_Sx SSAGA_Alc_D4_Dp_Sx SSAGA_Alc_Hvy_Drinks_Per_Day SSAGA_Alc_Hvy_Frq SSAGA_Alc_Hvy_Frq_5plus SSAGA_Alc_Hvy_Frq_Drk SSAGA_Alc_Hvy_Max_Drinks SSAGA_ChildhoodConduct SSAGA_Depressive_Ep SSAGA_Depressive_Sx SSAGA_Mj_Ab_Dep SSAGA_Mj_Times_Used SSAGA_Mj_Use SSAGA_PanicDisorder SSAGA_TB_Smoking_History SSAGA_TB_Still_Smoking SSAGA_Times_Used_Cocaine SSAGA_Times_Used_Hallucinogens SSAGA_Times_Used_Illicits SSAGA_Times_Used_Opiates SSAGA_Times_Used_Sedatives SSAGA_Times_Used_Stimulants Taste_AgeAdj Taste_Unadj THC Times_Used_Any_Tobacco_Today Total_Any_Tobacco_7days Total_Beer_Wine_Cooler_7days Total_Cigarettes_7days Total_Drinks_7days Total_Hard_Liquor_7days Total_Wine_7days VSPLOT_CRTE VSPLOT_OFF VSPLOT_TC. |

**Table S10.** The weights of behavioral measures and model-predicted thoughts in the primary identified CCA mode. **(a)** The set of behavioral measures with the strongest association ($r > .15$). Variables included in the CCA mode are marked in blue whereas variables whose weights are calculated post hoc are marked in gray. Variables are in their formal variable names defined here: https://wiki.humanconnectome.org/docs/assets/HCP_S1200_DataDictionary_Oct_30_2023.csv.
(**b**) The weights of model-predicted thoughts. Dimensions are marked in purple and topics are marked in green.

**a**

| *r* | %var | Variable name |
|---|---|---|
| 0.51 | 25.5 | SSAGA_TB_Yrs_Since_Quit |
| 0.40 | 16.0 | LifeSatisf_Unadj |
| 0.40 | 15.7 | EmotSupp_Unadj |
| 0.34 | 11.9 | SSAGA_Educ |
| 0.33 | 11.1 | PosAffect_Unadj |
| 0.33 | 10.8 | InstruSupp_Unadj |
| 0.32 | 10.4 | DDisc_AUC_40K |
| 0.31 | 9.6 | MeanPurp_Unadj |
| 0.31 | 9.4 | DDisc_SV_5yr_40K |
| 0.30 | 9.2 | NEOFAC_E |
| 0.30 | 9.0 | DDisc_AUC_200 |
| 0.30 | 8.9 | PicSeq_AgeAdj |
| 0.30 | 8.8 | DDisc_SV_3yr_40K |
| 0.30 | 8.8 | PicSeq_Unadj |
| 0.30 | 8.7 | PicVocab_Unadj |
| 0.29 | 8.7 | Friendship_Unadj |
| 0.29 | 8.5 | PicVocab_AgeAdj |
| 0.29 | 8.3 | DDisc_SV_3yr_200 |
| 0.29 | 8.2 | Endurance_AgeAdj |
| 0.29 | 8.1 | Endurance_Unadj |
| 0.28 | 8.0 | DDisc_SV_6mo_200 |
| 0.28 | 7.9 | DDisc_SV_1yr_200 |
| 0.28 | 7.7 | SSAGA_Income |
| 0.27 | 7.4 | NEOFAC_A |
| 0.27 | 7.1 | DDisc_SV_1yr_40K |
| 0.26 | 6.9 | DDisc_SV_10yr_40K |
| 0.26 | 6.6 | Dexterity_AgeAdj |
| 0.26 | 6.5 | Dexterity_Unadj |
| 0.25 | 6.4 | DDisc_SV_5yr_200 |
| -0.25 | 6.4 | SSAGA_Mj_Ab_Dep |
| -0.26 | 6.7 | SSAGA_Times_Used_Hallucinogens |
| -0.26 | 6.8 | ASR_Intn_Raw |
| -0.27 | 7.2 | ASR_Intn_T |
| -0.27 | 7.4 | ASR_Crit_Raw |
| -0.28 | 8.1 | ASR_Thot_Pct |
| -0.29 | 8.2 | ASR_Thot_Raw |
| -0.29 | 8.2 | SSAGA_FTND_Score |
| -0.30 | 8.7 | PercReject_Unadj |
| -0.30 | 8.9 | SSAGA_Times_Used_Sedatives |
| -0.31 | 9.6 | Loneliness_Unadj |
| -0.32 | 10.0 | SSAGA_Times_Used_Illicits |
| -0.33 | 10.7 | SSAGA_TB_Smoking_History |
| -0.33 | 10.8 | SSAGA_Mj_Times_Used |
| -0.35 | 12.2 | DSM_Antis_Raw |
| -0.37 | 13.6 | ASR_Witd_Raw |
| -0.37 | 13.7 | DSM_Antis_Pct |
| -0.37 | 13.8 | ASR_Witd_Pct |
| -0.40 | 15.9 | ASR_Rule_Pct |
| -0.41 | 17.2 | ASR_Rule_Raw |
| -0.48 | 22.8 | THC |
| -0.50 | 25.3 | SSAGA_TB_Still_Smoking |
| -0.51 | 26.2 | Times_Used_Any_Tobacco_Today |
| -0.55 | 30.3 | Avg_Weekend_Cigarettes_7days |
| -0.55 | 30.6 | Avg_Weekday_Cigarettes_7days |
| -0.57 | 32.0 | Avg_Weekday_Any_Tobacco_7days |
| -0.57 | 32.1 | Total_Cigarettes_7days |
| -0.57 | 32.4 | Avg_Weekend_Any_Tobacco_7days |
| -0.57 | 32.9 | Num_Days_Used_Any_Tobacco_7days |
| -0.58 | 34.1 | Total_Any_Tobacco_7days |

↑ Positive
↓ Negative

**Included in CCA**
Not included in CCA

**b**

| *r* | %var | Thoughts |
|---|---|---|
| 0.56 | 0.31 | Future planning |
| 0.43 | 0.18 | Other people |
| 0.35 | 0.12 | Social |
| 0.33 | 0.11 | Past memory |
| 0.33 | 0.11 | Awake |
| 0.27 | 0.07 | Future |
| 0.25 | 0.06 | Zoning out |
| 0.21 | 0.04 | Obligations |
| 0.02 | 0.00 | Image |
| 0.01 | 0.00 | Past |
| -0.07 | 0.00 | Detail |
| -0.10 | 0.01 | Valence |
| -0.16 | 0.03 | External |
| -0.33 | 0.11 | Current environment |
| -0.33 | 0.11 | Word |
| -0.68 | 0.46 | Body sensations and introspections |

↑ Positive
↓ Negative

**Thought dimensions** **Thought topics**

**Table S11.** Summary characteristics of null-thought models. We compared the number of selected edges and predictive accuracy of the real thought models against those obtained from the null models.

| Dimensions | Number of selected edges | | | | Predictive accuracy | | | |
|---|---|---|---|---|---|---|---|---|
| | real model | mean | 95% CI | *p*-value | real model | mean | 95% CI | *p*-value |
| Awake | 1024 | 919.2 | [728.6, 1182.2] | 0.182 | 0.077 | -0.0026 | [-0.101, 0.900] | 0.029* |
| External | 1300 | 908.4 | [728.8, 1197.8] | 0.003** | 0.104 | -0.0025 | [-0.100, 0.079] | 0.010* |
| Future | 1268 | 914.5 | [708.5, 1158.0] | 0.006** | 0.128 | -0.0016 | [-0.095, 0.085] | < 0.001*** |
| Past | 867 | 920.1 | [729.7, 1175.3] | 0.669 | -0.027 | 0.0034 | [-0.092, 0.080] | 0.753 |
| Other | 967 | 920.9 | [724.0, 1167.3] | 0.313 | 0.045 | 0.0000 | [-0.091, 0.091] | 0.175 |
| Valence | 1128 | 914.3 | [726.0, 1179.3] | 0.0502 | 0.112 | -0.0011 | [-0.010, 0.086] | 0.004** |
| Image | 1212 | 917.8 | [721.0, 1172.0] | 0.013* | 0.097 | -0.0004 | [-0.103, 0.089] | 0.011* |
| Word | 956 | 913.4 | [726.9, 1166.6] | 0.330 | 0.013 | -0.0033 | [-0.092, 0.086] | 0.362 |
| Detail | 759 | 915.8 | [723.8, 1178.9] | 0.932 | -0.031 | -0.0009 | [-0.095, 0.080] | 0.738 |

**Table S12.** Variants of the CCA analysis linking model-predicted thoughts with behavior. We observed statistically significant results in all versions of this analysis.

**CCA linking predicted thoughts and behavior**

| Models | *r*-values | corrected-*p* |
|---|---|---|
| 9 dimensional decoders + topic decoder (Fig. 5b) | 0.317 | $p < 0.001$*** |
| 5 significant dimensional decoders only<br>(Awake, External, Future, Valence, Image) | 0.238 | $p = 0.0387$* |
| Multidimensional decoder only | 0.301 | $p < 0.001$*** |
| Multidimensional decoder + control FD | 0.300 | $p < 0.001$*** |
| Multidimensional decoder + control pupil | 0.305 | $p < 0.001$*** |
| 9 dimensional decoders + topic decoder + control FD | 0.314 | $p < 0.001$*** |
| 9 dimensional decoders + topic decoder + control pupil | 0.312 | $p < 0.001$*** |
| 9 dimensional decoders only | 0.276 | $p = 0.0095$** |
| Topic decoder only | 0.280 | $p = 0.0071$** |
| Multidimensional decoder + topic decoder | 0.316 | $p < 0.001$*** |
| 5 significant dimensional decoders + topic decoder | 0.294 | $p = 0.0013$** |

**: $p < 0.01$, ***: $p < 0.001$
